# A large collection of novel nematode-infecting microsporidia and their diverse interactions with *C. elegans* and other related nematodes

**DOI:** 10.1101/074757

**Authors:** Gaotian Zhang, Martin Sachse, Marie-Christine Prevost, Robert Luallen, Emily Troemel, Marie-Anne Félix

**Affiliations:** Institut de Biologie de l’Ecole Normale Supérieure, CNRS, Inserm, ENS, PSL Research University, 75005 Paris, France.; School of Life Sciences, East China Normal University, 200062 Shanghai, China; Ultrapole, Institute Pasteur, 75015 Paris, France.; Division of Biological Sciences, Section of Cell and Developmental Biology, University of California San Diego, La Jolla, California, 92093 United States of America.

## Abstract

Microsporidia are fungi-related intracellular pathogens that may infect virtually all animals, but are poorly understood. The nematode *Caenorhabditis elegans* has recently become a model host for studying microsporidia through the identification of its natural microsporidian pathogen *Nematocida parisii.* However, it was unclear how widespread and diverse microsporidia infections are in *C. elegans* or other related nematodes in the wild. Here we describe the isolation and culture of 47 nematodes with microsporidian infections. *N. parisii* is found to be the most common microsporidia infecting *C. elegans* in the wild. In addition, we further describe and name six new species in the *Nematocida* genus. Our sampling and phylogenetic analysis further identify two subclades that are genetically distinct from *Nematocida*, and we name them *Enteropsectra* and *Pancytospora.* Interestingly, unlike *Nematocida,* these two genera belong to the main clade of microsporidia that includes human pathogens. All of these microsporidia are horizontally transmitted and most specifically infect intestinal cells, except *Pancytospora epiphaga* that replicates mostly in the epidermis of its *Caenorhabditis* host. At the subcellular level in the infected host cell, spores of the novel genus *Enteropsectra* show a characteristic apical distribution and exit via budding off of the plasma membrane, instead of exiting via exocytosis as spores of *Nematocida.* Host specificity is broad for some microsporidia, narrow for others: indeed, some microsporidia can infect *Oscheius tipulae* but not its sister species, and conversely. We also show that *N. ausubeli* fails to strongly induce in *C. elegans* the transcription of genes that are induced by other *Nematocida* species, suggesting it has evolved mechanisms to prevent induction of this host response. Altogether, these newly isolated species illustrate the diversity and ubiquity of microsporidian infections in nematodes, and provide a rich resource to investigate host-parasite coevolution in tractable nematode hosts.

**Author Summary:** Microsporidia are microbial parasites that live inside their host cells and can cause disease in humans and many other animals. The small nematode worm *Caenorhabditis elegans* has recently become a convenient model host for studying microsporidian infections. In this work, we sample *Caenorhabditis* and other small nematodes and 47 associated microsporidian strains from the wild. We characterize the parasites for their position in the evolutionary tree of microsporidia and for their lifecycle and morphology. We find several new species and genera, especially some that are distantly related to the previously known *Nematocida parisii* and instead closely related to human pathogens. We find that some of these species have a narrow host range. We studied two species in detail using electron microscopy and uncover a new likely mode of exit from the host cell, by budding off the host cell plasma membrane rather than by fusion of a vesicle to the plasma membrane as in *N. parisii.* We also find a new species that infects the epidermis and muscles of *Caenorhabditis* rather than the host intestinal cells and is closely related to human pathogens. Finally, we find that one *Nematocida* species fails to elicit the same host response that other *Nematocida* species do. These new microsporidia open up many windows into microsporidia biology and opportunities to investigate host-parasite coevolution in the *C. elegans* system.

## INTRODUCTION

Microsporidia are fungi-related obligate intracellular pathogens, with over 1400 described species [1,2]. Interest in these organisms started 150 years ago when researchers, especially Louis Pasteur, studied silkworm disease that was caused by a microsporidian species later named *Nosema bombycis* [3]. In the past decades, microsporidia have attracted more attention when they were revealed to be a cause of diarrhea in immunocompromised patients and were further demonstrated to have a high prevalence in some areas in immunocompetent patients and healthy individuals [4–6].

Microsporidia are transmitted between hosts through a spore stage. Inside the microsporidian spore is found a characteristic structure called the polar tube, which at the time of infection can pierce through host cell membranes and introduce the sporoplasm (the spore cytoplasm and nucleus) into host cells [1,7]. These obligate intracellular pathogens are known to infect a wide range of hosts among protists and animals, especially insects, fish and mammals [1]. Even though nematodes constitute a huge phylum with over 25,000 described species, very few studies on microsporidian infections in nematodes have been reported so far [1].

The free-living nematode *Caenorhabditis elegans* has been used as a major biological model species over the last 50 years [8]. However, until the past decade, little was known about its biology and ecology in its natural environment and no natural pathogens were isolated until *C. elegans* could be readily isolated from natural environments. *C. elegans* is now known to be found in compost heaps, rotting fruits (apples, figs, etc.) and herbaceous stems, as well as with diverse carrier invertebrates (snails, isopods, etc.) [9–11]. *C. elegans* inhabits a community with a variety of prokaryotic and eukaryotic microbes, including both its food and pathogens, which likely have a large impact on its physiology and evolution [12–15].

With an improved understanding of the natural history of *Caenorhabditis* [16,17], dramatically increased number of various wild rhabditid nematode strains and species have been isolated and identified. *C. elegans*’ close relatives such as *Caenorhabditis briggsae* or *Caenorhabditis remanei* are isolated from similar environments [18]. *Oscheius tipulae,* a very common bacteriovorous nematode species, also in family Rhabditidae, can be readily isolated from soil and rotting vegetal matter [10,19,20], as well as its closest known relative is *Oscheius* sp. 3, with which it cannot interbreed [21]. Interest in these rhabditid nematodes concerns not only the evolution of genomic and phenotypic characters, but also their inter-and intraspecific interactions and co-evolution with other organisms, especially with various microbes in their natural habitats. While nematodes feed on bacteria and small eukaryotes, some microbes take nematodes as their food source [13,14,16]. Among them, microsporidia are obligate intracellular parasites and thus in particularly tight association with their hosts.

The microsporidian *Nematocida parisii* was the first found natural intracellular pathogen of *C. elegans,* which we isolated from a wild *C. elegans* sampled near Paris, France [22]. *Nematocida* sp. 1 (described here as *Nematocida ausubeli*) was further isolated from a wild *Caenorhabditis briggsae* strain in India [22]. A microsporidian species isolated in *C. elegans* was found to infect the epidermis and muscles and was named *Nematocida displodere* [23]. Two microsporidia infecting marine nematodes have also been described, namely *Sporanauta perivermis* [24] and *Nematocenator marisprofundi* [25,26]. However, the extent and diversity of microsporidia infections in nematodes remained sparsely described.

Here we describe a collection of 47 terrestrial nematode strains that we isolated from the wild with a microsporidian infection. The microsporidia can be grown in the laboratory in their host using *C. elegans* culture conditions and stored frozen with their nematode host. They are all transmitted horizontally. In this set, we found that *N. parisii* and *N. ausubeli* (formerly called *N.* sp. 1) are in association with further host species and display a wider geographical distribution than so far reported [22]. *N. parisii* is the most common *C.* elegans-infecting species we found in the wild. We further discovered new nematode-infecting microsporidian species. From our phylogenetic analysis using small subunit (SSU) ribosomal DNA and β-tubulin sequences, five new microsporidia species were placed in the *Nematocida* genus, while the others defined two new genera in the microsporidian clade often designated as Clade IV, which includes human pathogenic microsporidia such as *Enterocytozoon bieneusi* and *Vittaforma corneae.* The similarities and differences in the morphological features of these microsporidia matched their groupings by sequence similarity. We therefore describe two new microsporidian genera, *Enteropsectra* and *Pancytospora.* and nine new species in these two genera and *Nematocida.* We further examined *Nematocida ausubeli* and *Enteropsectra longa* by electron microscopy, which allowed us to observe different mechanisms for their exit from host intestinal cells, through a vesicular pathway for *N. ausubeli* (as described for *N. parisii*; [27]), but surprisingly through membrane budding for *E. longa.* Concerning specificity of infection, we find cases of tight specificity between host and pathogen. We also find that *N. ausubeli* fails to strongly induce the transcription of genes that are induced in *C. elegans* by *N. parisii* infection. Overall, our study points to strong and diverse interactions between wild rhabditid nematodes and microsporidia, and provides a platform for further study of these infections.

## RESULTS

### A large collection of microsporidian-infected nematode cultures

Our worldwide sampling of bacteriovorous nematodes was primarily aimed at isolating *Caenorhabditis* species and, to a lesser degree, *Oscheius* species. From this sampling, we identified a subset of strains with a pale body color (Fig 1A), some of which, upon morphological examination using Nomarski microscopy [22], appeared infected with microsporidia. In total, we collected 47 nematode strains (S1 Table) displaying putative microsporidian infections, comprising 10 nematode species from different parts of the world (Table 1, 2; Fig 1B). The microsporidia strain JUm2807 was isolated during these sampling efforts and described elsewhere as *Nematocida displodere,* and is not considered here [23].

**Table 1:** Collection of wild nematode-infecting microsporidia strains.

*Nematocida parisii* infection
| Host | Strain | Country / Year | Sample composition | Spore feature |
| --- | --- | --- | --- | --- |
| *Caenorhabditis elegans* | **ERTm1**<sup>a</sup> | France / 2004 | compost | 2S<sup>c</sup> |
|  | ERTm3<sup>a</sup> | France / 2007 | rotting apple | 2S<sup>c</sup> |
|  | JUm1248<sup>a b</sup> | France / 2007 | rotting apple | n. d. |
|  | JUm1249<sup>b</sup> | France / 2007 | rotting apple | 2S |
|  | JUm1253<sup>b</sup> | France / 2007 | rotting apple | n. d. |
|  | ERTm4<sup>a</sup> | France / 2008 | compost | 2S<sup>c</sup> |
|  | JUm1762<sup>b</sup> | France / 2009 | rotting stem | 2S |
|  | JUm1893<sup>b</sup> | France / 2009 | rotting apple | 2S |
|  | JUm2106 | France / 2011 | compost | 2S |
|  | JUm2131 | France / 2011 | rotting stem | 2S |
|  | JUm2132 | France / 2011 | rotting stem | 2S |
|  | JUm2287 | France / 2013 | compost | 2S |
|  | JUm2816 | France / 2014 | compost | n. d. |
| *Caenorhabditis briggsae* | JUm1254 | France / 2007 | rotting apple | 2S |
|  | ERTm5<sup>a</sup> | Hawaii / 2011 | rotting fruit | 2S |
|  | JUm2590 | France / 2013 | rotting apple | 2S |
|  | JUm2793 | France / 2014 | compost | n. d. |

**B. *Nematocida ausubeli* (*N. sp.* 1) infection**
| Host | Strain | Country / Year | Sample composition | Spore feature |
| --- | --- | --- | --- | --- |
| *Caenorhabditis elegans* | JUm2009 | France / 2010 | rotting apples | 2S |
|  | JUm2520 | Portugal / 2013 | rotting fruit | 2S |
|  | JUm2526 | Portugal / 2013 | rotting fruit | 2S |
|  | JUm2586 | France / 2013 | rotting pear | 2S |
|  | JUm2671 | France / 2011 | rotting apple | 2S |
|  | JUm2825 | France / 2014 | rotting stem | 2S |
| *Caenorhabditis briggsae* | **ERTm2**<sup>a</sup> | India / 2007 | rotting mix | 2S<sup>c</sup> |
|  | ERTm6<sup>a</sup> | Cape Verde / 2009 | soil | 2S |
|  | JUm2799 | Germany / 2014 | rotting fruits | 2S |
| *Caenorhabditis remanei* | JUm2796 | Germany / 2014 | rotting stems | n. d. |

The reference strain of each newly found species is in bold. 2S: two distinct sizes of spores; n. d.: not determined. <sup>a</sup> Strains and names published previously; <sup>b</sup> 18S sequences courtesy of Aurore Dubuffet and Hinrich Schulenburg; <sup>c</sup> Morphological feature described in Troemel et al. 2008.

**Table 2:** Collection of other microsporidia species infecting wild nematodes.

| Microsporidia | Nematode host | Strain | Country / Year | Sample composition | Spore feature |
| --- | --- | --- | --- | --- | --- |
| *Nematocida* | *C. briggsae* | **JUm2507** | Thailand / 2013 | rotting fruit | 2S |
| *major* n. sp. | *C. briggsae* | JUm2747 | Guadeloupe / 2014 | rotting fruit | 2S |
|  | *C. tropicalis* | JUm2751 | Guadeloupe / 2014 | rotting fruit | n. d. |
| *Nematocida* | *Oscheius tipulae* | **JUm1510** | Czech Republic / 2008 | rotting apple | 2S |
| *minor* n. sp. | *O. tipulae* | JUm2772 | Armenia / 2014 | rotting fruit | n. d. |
| *Nematocida* | *O. tipulae* | **JUm1504** | France / 2008 | rotting stem | 1S |
| *homosporus* n. sp. | *Rhabditella typhae* | NICm516 | Portugal / 2013 | rotting apple | 1S |
| *Nematocid* | *Procephalobus* sp. | **JUm2895** | Spain / 2015 | rotting fruits | 1S |
| *ciargi* n. sp. |  |  |  |  |  |
| *Nematocida* | *C. sp. 42* | **NICm1041** | French Guiana / 2014 | flower | n. d. |
| sp. 7 |  |  |  |  |  |
| *Enteropsectra* | *Oscheius* sp. 3 | **JUm408** | Iceland / 2002 | compost | 1S, LT, AP |
| *longa* n. sp. |  |  |  |  |  |
| *Enteropsectra* | *O. tipulae* | **JUm2551** | France / 2013 | rotting apple | 1S, SR, AP |
| *breve* n. sp. | *O. tipulae* | JUm1483 | France / 2008 | rotting plum | 1S, SR, AP |
|  | *O. tipulae* | JUm1456 | France / 2008 | rotting fruit | n. d. |
| *Pancytospora* | *O. tipulae* | **JUm1505** | France / 2008 | rotting apple | LT |
| *philotis* n. sp. | *O. tipulae* | JUm1460 | France / 2008 | rotting snail | LT |
|  | *O. tipulae* | JUm1670 | France / 2009 | rotting apple | LT |
|  | *O. tipulae* | JUm2552 | France / 2013 | rotting stem | LT |
| *Pancytospora* | *C. brenneri* | **JUm1396** | Colombia / 2008 | rotting fruit | LT |
| *epiphaga* n. sp. |  |  |  |  |  |

**Fig 1:**
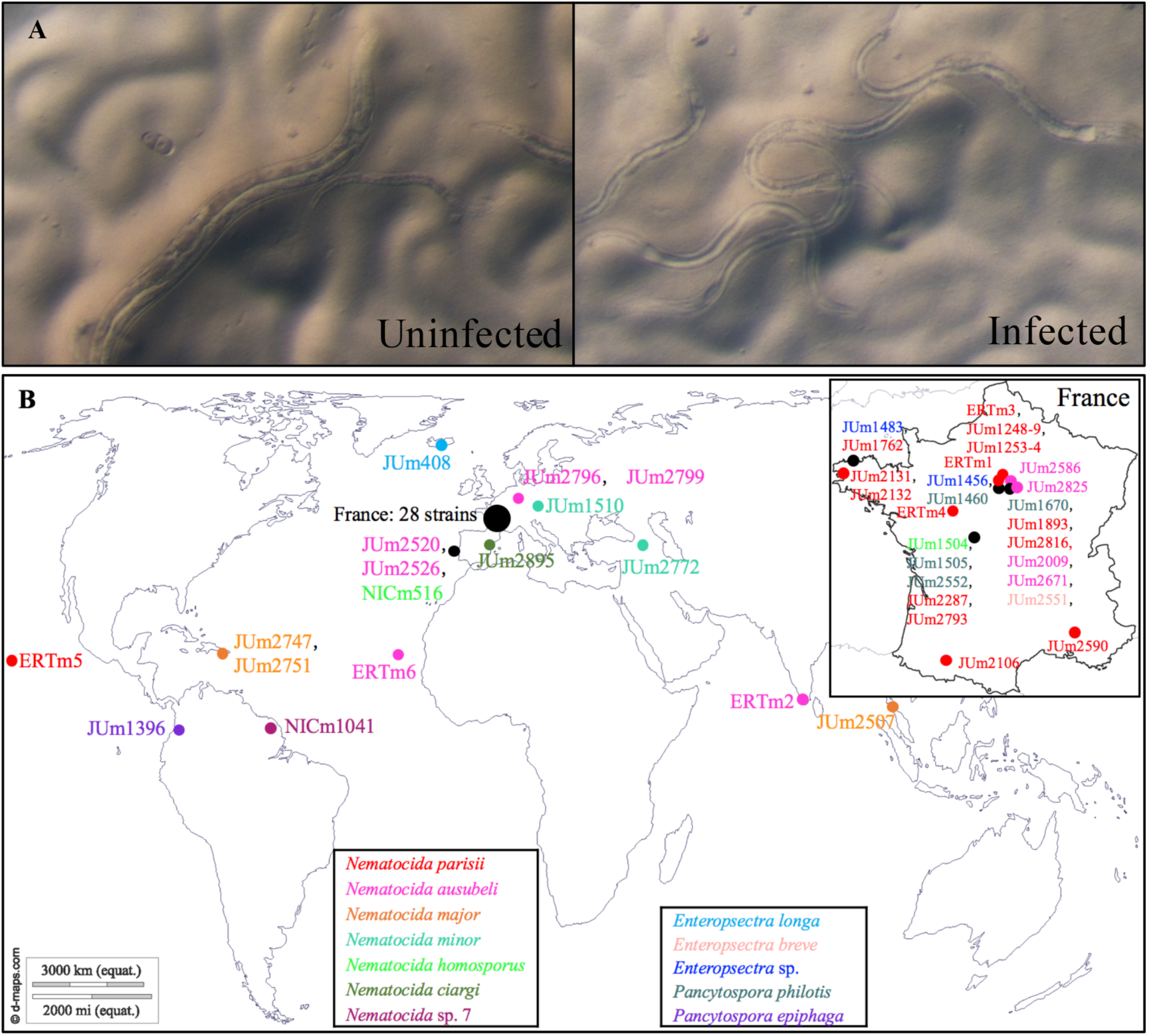
Isolation of nematode-infecting microsporidia. A. Morphological screen for infected worms (here *C. elegans* JU2009). Compared to uninfected worms, infected adult worms have a paler body color and are generally smaller and thinner. Here the intestinal cells are reduced in width, which often occurs in late infections (see also S1F,G Fig). Note that the pale body color may result from many environmental conditions, and thus these animals were further screened by Nomarski optics for microsporidian infections. B. Geographic distribution of our collection of nematode-infecting microsporidia. Sampling locations are represented by differently colored symbols based on microsporidian species. Black symbols were used when different microsporidian species were found in the same location.

**Fig 2:**
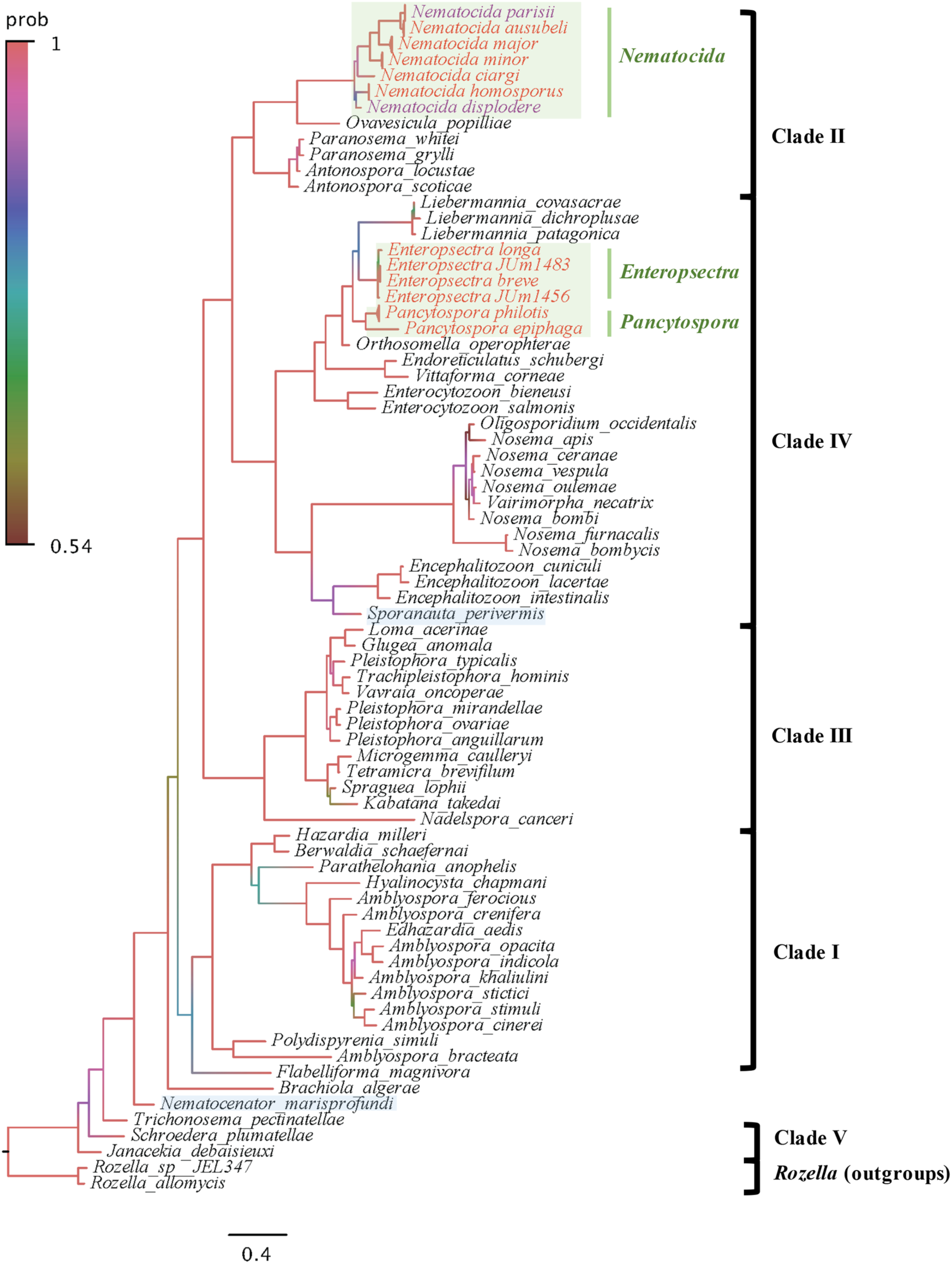
Bayesian inference SSU rDNA phylogeny of microsporidia species. SSU rDNA sequences from 45 nematode-infecting microsporidia species and 60 other microsporidia species in the databases were used. The tree was generated using MrBayes v3.2.2 and refined by FigTree v1.4.2. Model Kimura 2-Parameter (K2P) was applied. Branch colors show the posterior probability, with the corresponding color code shown on the left. The light green boxes designate microsporidia infecting terrestrial nematodes and light-blue rectangles designate those infecting marine nematodes. Scale bar indicates expected changes per site. Branches of species with more than one strain were compressed.

The unidentified microsporidian strains were characterized by sequencing of PCR fragments of the SSU rDNA and β-tubulin genes. We were able to amplify 45 SSU rDNA sequences (most 1390 bp long) and 32 β-tubulin sequences (most 1210 bp long) (S1 Table). We first blasted the sequences in GenBank for initial grouping, then built phylogenetic trees and calculated interspecific genetic distances, based on our sequences and the sequences of related species from GenBank. We present below the grouping and phylogenetic distribution of new microsporidia strains, starting with those closest to *N. parisii.*

### *N. parisii* and *N. ausubeli* are commonly found in *Caenorhabditis* nematodes

Molecular sequences of microsporidia in ten wild *C. elegans* strains and four *C. briggsae* strains showed ≥ 99% SSU rDNA and ≥ 97% β-tubulin sequence identities to *N. parisii* sequences in GenBank. In the global phylogenetic analysis of microsporidia, these 14 sequences form a group with previously reported sequences of *N. parisii* strains ERTm1, ERTm3 and ERTm5 [26] (Fig 2). The *N. parisii* isolates were all found in Europe (note however that the sampling is highly biased towards Europe, especially France), with the exception of the previously reported ERTm5 (JUm2055), isolated from a *C. briggsae* strain sampled in Hawaii (Fig 1B; Table 1A) [28].

Eight other microsporidian strains showed ≥ 99% SSU rDNA and ≥ 95% β-tubulin sequence identities to the corresponding genes of the unnamed *Nematocida* sp. 1 in GenBank (Table 1B), previously reported in *C. briggsae* [22,29]. This N. sp. 1 group is most closely related to *N. parisii* in the microsporidian phylogeny and the sequences of both SSU and β-tubulin genes gave the same grouping (Fig 2, 3; S2 Fig; Table 3). Because of these new samples of N. sp. 1 and their phylogenetic difference and genetic distance to the *N. parisii* group, here we describe *N*. sp. 1 as *Nematocida ausubeli* n. sp. (see Taxonomy section after the Discussion). Whereas N. *ausubeli* was so far only reported from *C. briggsae* (India, Cape Verde [29]), we also found it in *C. elegans* and *C. remanei*, in France, Portugal and Germany (Table 1B; Fig 1B), thus broadening its geographic and host range to several species of the *Elegans* group of *Caenorhabditis* from Europe.

**Table 3:**
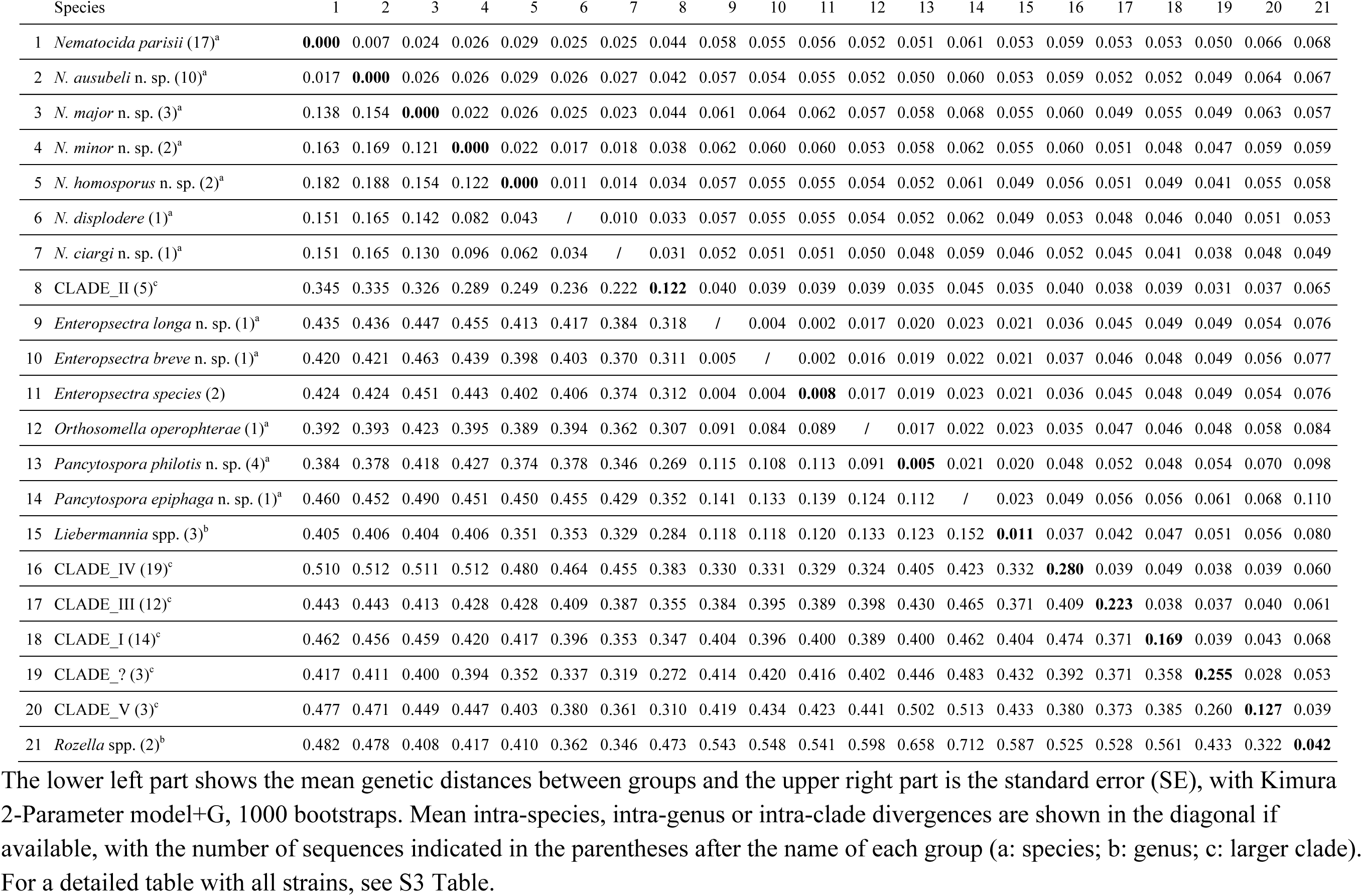
Molecular distances of microsporidia SSU rDNA.

| Species | 1 | 2 | 3 | 4 | 5 | 6 | 7 | 8 | 9 | 10 | 11 | 12 | 13 | 14 | 15 | 16 | 17 | 18 | 19 | 20 | 21 |
| --- | --- | --- | --- | --- | --- | --- | --- | --- | --- | --- | --- | --- | --- | --- | --- | --- | --- | --- | --- | --- | --- |
| 1 *Nematocida parisii* (17)<sup>a</sup> | **0.000** | 0.007 | 0.024 | 0.026 | 0.029 | 0.025 | 0.025 | 0.044 | 0.058 | 0.055 | 0.056 | 0.052 | 0.051 | 0.061 | 0.053 | 0.059 | 0.053 | 0.053 | 0.050 | 0.066 | 0.068 |
| 2 *N. ausubeli* n. sp. (10)<sup>a</sup> | 0.017 | **0.000** | 0.026 | 0.026 | 0.029 | 0.026 | 0.027 | 0.042 | 0.057 | 0.054 | 0.055 | 0.052 | 0.050 | 0.060 | 0.053 | 0.059 | 0.052 | 0.052 | 0.049 | 0.064 | 0.067 |
| 3 *N. major* n. sp. (3)<sup>a</sup> | 0.138 | 0.154 | **0.000** | 0.022 | 0.026 | 0.025 | 0.023 | 0.044 | 0.061 | 0.064 | 0.062 | 0.057 | 0.058 | 0.068 | 0.055 | 0.060 | 0.049 | 0.055 | 0.049 | 0.063 | 0.057 |
| 4 *N. minor* n. sp. (2)<sup>a</sup> | 0.163 | 0.169 | 0.121 | **0.000** | 0.022 | 0.017 | 0.018 | 0.038 | 0.062 | 0.060 | 0.060 | 0.053 | 0.058 | 0.062 | 0.055 | 0.060 | 0.051 | 0.048 | 0.047 | 0.059 | 0.059 |
| 5 *N. homosporus* n. sp. (2)<sup>a</sup> | 0.182 | 0.188 | 0.154 | 0.122 | **0.000** | 0.011 | 0.014 | 0.034 | 0.057 | 0.055 | 0.055 | 0.054 | 0.052 | 0.061 | 0.049 | 0.056 | 0.051 | 0.049 | 0.041 | 0.055 | 0.058 |
| 6 *N. displodere* (1)<sup>a</sup> | 0.151 | 0.165 | 0.142 | 0.082 | 0.043 | / | 0.010 | 0.033 | 0.057 | 0.055 | 0.055 | 0.054 | 0.052 | 0.062 | 0.049 | 0.053 | 0.048 | 0.046 | 0.040 | 0.051 | 0.053 |
| 7 *N. ciargi* n. sp. (1)<sup>a</sup> | 0.151 | 0.165 | 0.130 | 0.096 | 0.062 | 0.034 | / | 0.031 | 0.052 | 0.051 | 0.051 | 0.050 | 0.048 | 0.059 | 0.046 | 0.052 | 0.045 | 0.041 | 0.038 | 0.048 | 0.049 |
| 8 CLADE_II (5)<sup>c</sup> | 0.345 | 0.335 | 0.326 | 0.289 | 0.249 | 0.236 | 0.222 | **0.122** | 0.040 | 0.039 | 0.039 | 0.039 | 0.035 | 0.045 | 0.035 | 0.040 | 0.038 | 0.039 | 0.031 | 0.037 | 0.065 |
| 9 *Enteropsectra longa* n. sp. (1)<sup>a</sup> | 0.435 | 0.436 | 0.447 | 0.455 | 0.413 | 0.417 | 0.384 | 0.318 | / | 0.004 | 0.002 | 0.017 | 0.020 | 0.023 | 0.021 | 0.036 | 0.045 | 0.049 | 0.049 | 0.054 | 0.076 |
| 10 *Enteropsectra breve* n. sp. (1)<sup>a</sup> | 0.420 | 0.421 | 0.463 | 0.439 | 0.398 | 0.403 | 0.370 | 0.311 | 0.005 | / | 0.002 | 0.016 | 0.019 | 0.022 | 0.021 | 0.037 | 0.046 | 0.048 | 0.049 | 0.056 | 0.077 |
| 11 *Enteropsectra species* (2) | 0.424 | 0.424 | 0.451 | 0.443 | 0.402 | 0.406 | 0.374 | 0.312 | 0.004 | 0.004 | **0.008** | 0.017 | 0.019 | 0.023 | 0.021 | 0.036 | 0.045 | 0.048 | 0.049 | 0.054 | 0.076 |
| 12 *Orthosomella operophterae* (1)<sup>a</sup> | 0.392 | 0.393 | 0.423 | 0.395 | 0.389 | 0.394 | 0.362 | 0.307 | 0.091 | 0.084 | 0.089 | / | 0.017 | 0.022 | 0.023 | 0.035 | 0.047 | 0.046 | 0.048 | 0.058 | 0.084 |
| 13 *Pancytospora philotis* n. sp. (4)<sup>a</sup> | 0.384 | 0.378 | 0.418 | 0.427 | 0.374 | 0.378 | 0.346 | 0.269 | 0.115 | 0.108 | 0.113 | 0.091 | **0.005** | 0.021 | 0.020 | 0.048 | 0.052 | 0.048 | 0.054 | 0.070 | 0.098 |
| 14 *Pancytospora epiphaga* n. sp. (1)<sup>a</sup> | 0.460 | 0.452 | 0.490 | 0.451 | 0.450 | 0.455 | 0.429 | 0.352 | 0.141 | 0.133 | 0.139 | 0.124 | 0.112 | / | 0.023 | 0.049 | 0.056 | 0.056 | 0.061 | 0.068 | 0.110 |
| 15 *Liebermannia* spp. (3)<sup>b</sup> | 0.405 | 0.406 | 0.404 | 0.406 | 0.351 | 0.353 | 0.329 | 0.284 | 0.118 | 0.118 | 0.120 | 0.133 | 0.123 | 0.152 | **0.011** | 0.037 | 0.042 | 0.047 | 0.051 | 0.056 | 0.080 |
| 16 CLADE_IV (19)<sup>c</sup> | 0.510 | 0.512 | 0.511 | 0.512 | 0.480 | 0.464 | 0.455 | 0.383 | 0.330 | 0.331 | 0.329 | 0.324 | 0.405 | 0.423 | 0.332 | **0.280** | 0.039 | 0.049 | 0.038 | 0.039 | 0.060 |
| 17 CLADE_III (12)<sup>c</sup> | 0.443 | 0.443 | 0.413 | 0.428 | 0.428 | 0.409 | 0.387 | 0.355 | 0.384 | 0.395 | 0.389 | 0.398 | 0.430 | 0.465 | 0.371 | 0.409 | **0.223** | 0.038 | 0.037 | 0.040 | 0.061 |
| 18 CLADE_I (14)<sup>c</sup> | 0.462 | 0.456 | 0.459 | 0.420 | 0.417 | 0.396 | 0.353 | 0.347 | 0.404 | 0.396 | 0.400 | 0.389 | 0.400 | 0.462 | 0.404 | 0.474 | 0.371 | **0.169** | 0.039 | 0.043 | 0.068 |
| 19 CLADE_? (3)<sup>c</sup> | 0.417 | 0.411 | 0.400 | 0.394 | 0.352 | 0.337 | 0.319 | 0.272 | 0.414 | 0.420 | 0.416 | 0.402 | 0.446 | 0.483 | 0.432 | 0.392 | 0.371 | 0.358 | **0.255** | 0.028 | 0.053 |
| 20 CLADE_V (3)<sup>c</sup> | 0.477 | 0.471 | 0.449 | 0.447 | 0.403 | 0.380 | 0.361 | 0.310 | 0.419 | 0.434 | 0.423 | 0.441 | 0.502 | 0.513 | 0.433 | 0.380 | 0.373 | 0.385 | 0.260 | **0.127** | 0.039 |
| 21 *Rozella* spp. (2)<sup>b</sup> | 0.482 | 0.478 | 0.408 | 0.417 | 0.410 | 0.362 | 0.346 | 0.473 | 0.543 | 0.548 | 0.541 | 0.598 | 0.658 | 0.712 | 0.587 | 0.525 | 0.528 | 0.561 | 0.433 | 0.322 | **0.042** |

The lower left part shows the mean genetic distances between groups and the upper right part is the standard error (SE), with Kimura 2-Parameter model+G, 1000 bootstraps. Mean intra-species, intra-genus or intra-clade divergences are shown in the diagonal if available, with the number of sequences indicated in the parentheses after the name of each group (a: species; b: genus; c: larger clade). For a detailed table with all strains, see S3 Table.

The remaining 20 microsporidia strains that we identified are distributed among several other species, including some species in another clade (see below). Thus the dominant microsporidia species in our collection of *Caenorhabditis* and other nematodes are *Nematocida parisii* and *Nematocida ausubeli* n. sp., with 17 and 10 strains, respectively. They were found in several species of the *Elegans* group of *Caenorhabditis* nematodes.

### Diversity of *Nematocida* species

Of the remaining 19 microsporidian strains, nine had a *Nematocida* species as their top blast hit in GenBank, with similarity between 81% ~ 86% of SSU rDNA and 74% ~ 84% of β-tubulin genes. In terms of host and geographical ranges, these microsporidia were found in two *C. briggsae* strains (Thailand and Guadeloupe), one *C. tropicalis* strain (Guadeloupe), one C. sp. 42 strain (NIC1041 from French Guiana), three *Oscheius tipulae* strains (France, Czech Republic, and Armenia), one *Rhabditella typhae* strain (Portugal) and one *Procephalobus* sp. strain (JU2895 from Spain). In the phylogenetic analysis of SSU rDNA, the corresponding sequences formed a single clade with *N. parisii* and *N. ausubeli*, with *Ovavesicula popilliae* as sister group within Clade II of the microsporidian phylum (see Fig 2) [33]. In addition, the JUm2807 strain that has been recently described as *Nematocida displodere* [23] is distinct from all of them.

From phylogenetic analysis and genetic distance of SSU rDNA genes, these *Nematocida* strains form four groups. These putative new *Nematocida* species have a mean genetic distance among them of at least 0.06 (Table 3), while their intra-specific genetic distances are all 0.00 (when several strains were isolated). This inter-group distance is also greater than the distance between *N. parisii* and *N. ausubeli*. Hence we describe them below as four new species: *Nematocida minor, Nematocida major, Nematocida homosporus* and *Nematocida ciargi* n. spp. (see Taxonomy section).

In terms of the phylogenetic relationships within the *Nematocida* genus in the SSU rDNA tree, the first outgroup clade to *N. parisii* + *N. ausubeli* was formed by JUm2751, JUm2747 and JUm2751, corresponding to *N. major* (Fig 2). The second branch out was formed by JUm1510 and JUm2772, described here as *N. minor. N. ciargi* JUm2895 was placed in a basal position to the clade formed by *N. parisii*, *N. ausubeli*, *N. major* and *N. minor* (Fig 2). At the base of the *Nematocida* genus, the most externally branching sequences appeared to be those of *N. displodere* JUm2807, and of *N. homosporus* JUm1504 and NICm516. All topologies were highly supported, except for the node defining the latter clade of *N. homosporus* and *N. displodere* (Fig 2). In the phylogenetic tree based on both genes (SSU rDNA and β-tubulin), *N. ciargi* was placed at the base of *Nematocida* genus, while *N. displodere* and *N. homosporus* still formed one clade (Fig 3). The phylogenetic tree only based on β-tubulin sequences supported the grouping of strains and overall their relative positions (S2 Fig), except that the relative placement of *N. displodere* and *N. ciargi* was exchanged. The β-tubulin phylogeny has one more branch formed by NICm1041, numbered provisionally *N.* sp. 7, for which we failed to amplify the SSU rDNA fragment. Whole-genome analysis could be performed in the future to refine these placements.

**Fig 3:**
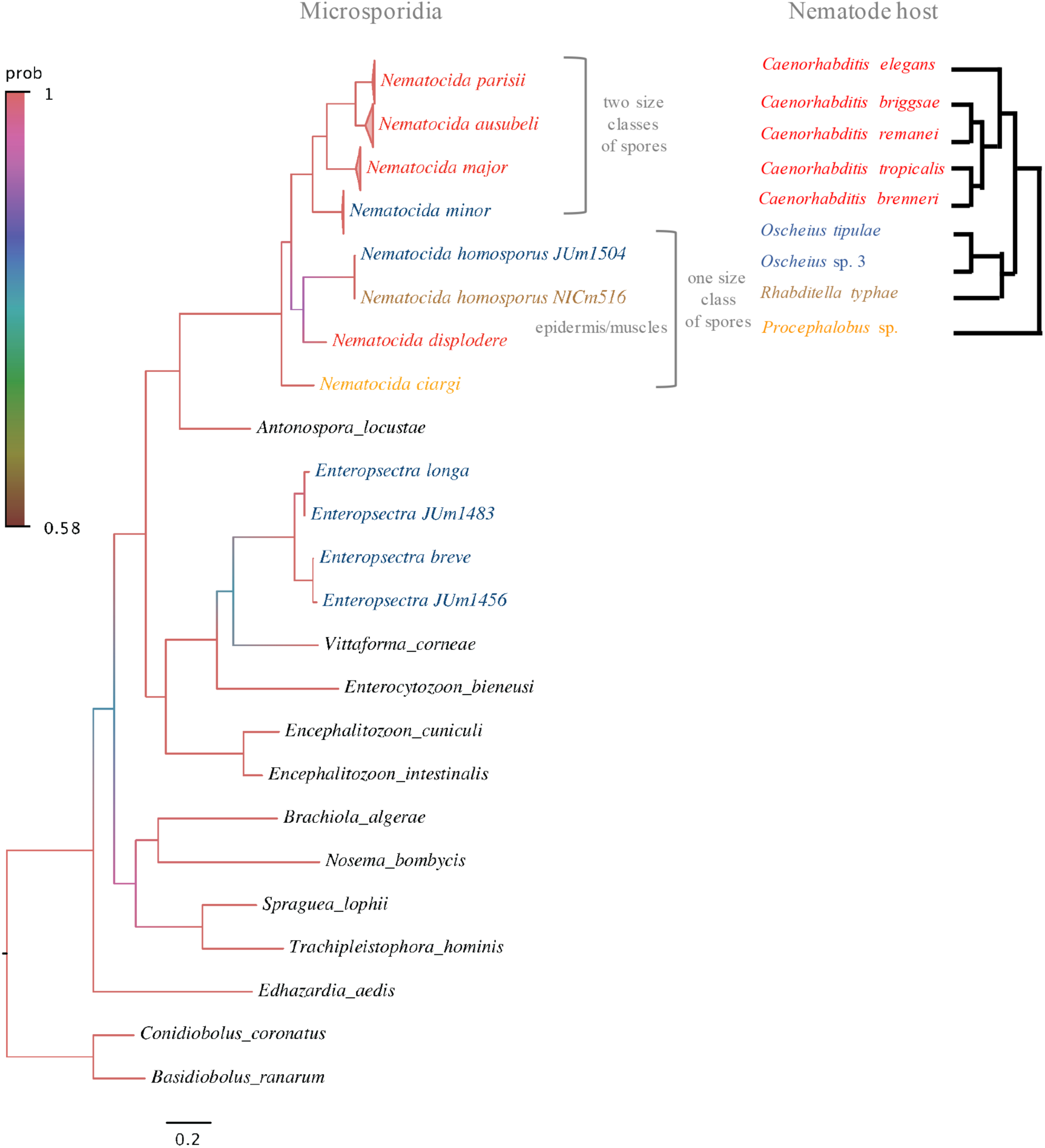
Bayesian inference phylogeny of concatenated SSU rDNA and β-tubulin sequences of 22 microsporidia species, and comparison with the nematode host phylogeny. Bayesian inference phylogeny (left) based on 49 sequences concatenated from SSU rDNA and β-tubulin genes of 22 microsporidia species. Model General time reversible (GTR) was applied. The branches were colored and annotated as in Fig 2. On the right is a diagram (generated based on phylogenies from [18,21,30–32]) showing the relative position of nematode species found with microsporidia infections. Nematode-infecting microsporidia pathogens and their hosts were colored based on host genus. Correspondent positions of nematode-infecting microsporidia and nematodes on their phylogenies indicate a possible coevolution of nematodes and their natural pathogenic microsporidia.

The *Nematocida* consensus phylogeny is shown in Fig 3 next to the phylogenetic relationships of the nematode hosts in which they were naturally found (see below for further specificity tests). Although the numbers of samples and species are too low for rigorous testing, the data are at least consistent with the intestinal microsporidia species branching through continuous co-evolution with their nematode host. For example, all intestinal *Nematocida* species found in *Caenorhabditis* species form one clade, with a first outgroup including *Oscheius* and *Rhabditella* pathogens and a distant outgroup infecting the distant outgroup *Procephalobus* (Fig 3). The exception is *N. displodere* that was found a single time, in *C. elegans,* and corresponds to a change in tissue tropism.

### Lifecycle of new *Nematocida* species

As with previously isolated *Nematocida,* the newly identified microsporidia appeared to be transmitted horizontally, because a bleaching treatment [34] of infected gravid adults eliminated the infection in the culture and reinfection could be obtained by exposure from spores in the environment. All *Nematocida* microsporidia stages described here were found exclusively in the intestinal cells and were not detected in the germ line.

As previously described for *N. parisii* [22], two main stages could be distinguished by Nomarski optics. First, the meront stage appeared as areas of infected intestinal cells devoid of storage granules. These areas were first small circular regions, then extending to longer grooves. Second, rod-shaped sporoblasts and spores appeared in the intestinal cell cytoplasm. In host cells that were heavily infected with *N. parisii* and some other species, groups of spores inside vesicles could be seen [22], possibly derived from spore re-endocytosis [35]. In this study, as described before [22], all *N. parisii* and *N. ausubeli* infections displayed two distinct classes of spore size (Table 1; Fig 4A, B; S3A Fig).

**Fig 4:**
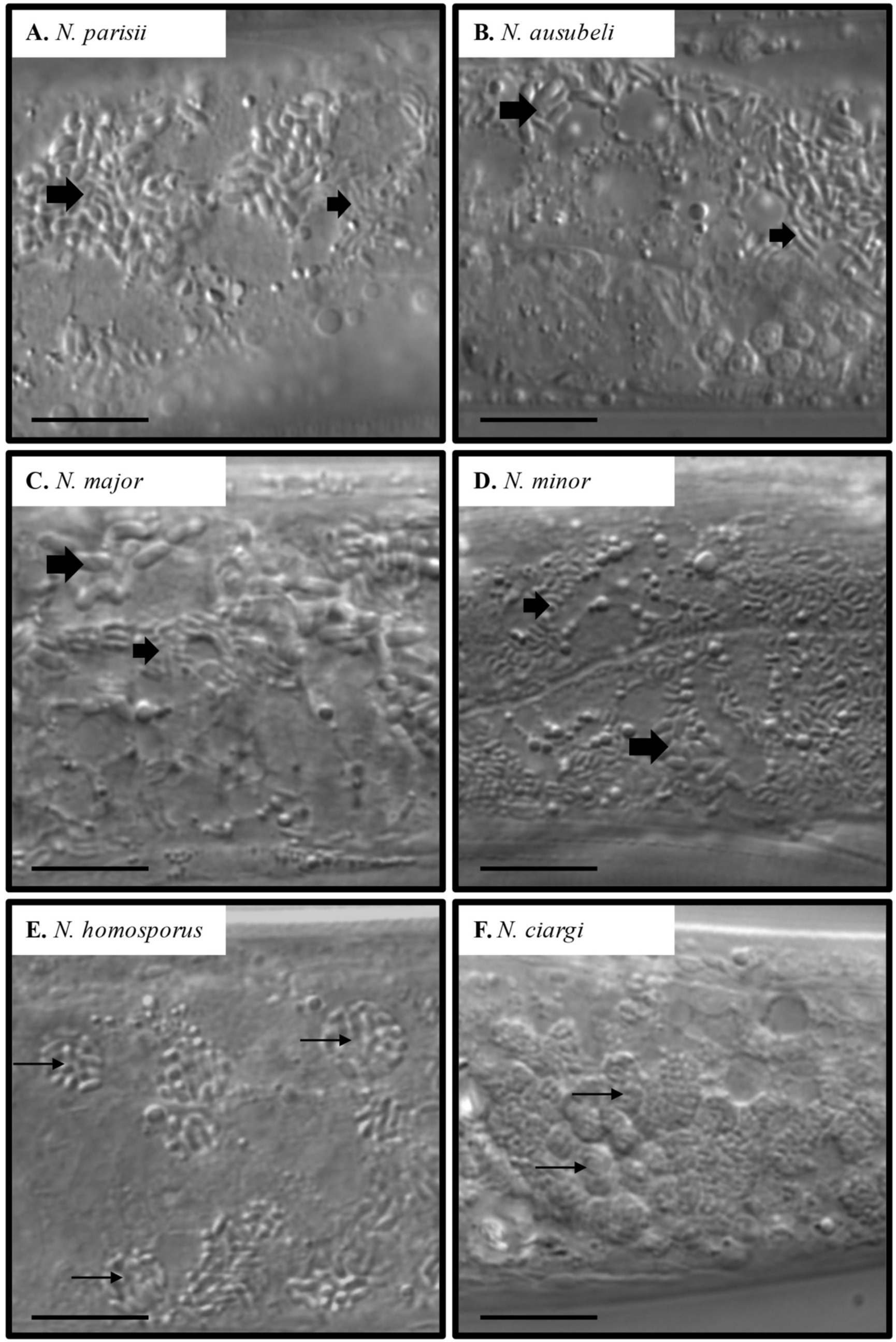
Spore morphology of the different *Nematocida* species by Nomarski optics. A. Wild *Caenorhabditis elegans* strain JU1249, with *Nematocida parisii* infection. **B.** Wild *C. elegans* strain JU2520, with *Nematocida ausubeli* infection. **C.** Wild **C.** *briggsae* stain JU2747, with *N. major* infection. D. Wild *Oscheius tipulae* strain JU1510, with *N. minor* infection. A ~ D, large and small spore classes are indicated by larger and smaller arrows, respectively. Spores in each class are smaller than those in A-C in the corresponding class. E. Wild *Rhabditella typhae* strain NIC516, with *N. homosporus* infection. A single class of spore size is observed, often clustered inside vesicles as indicated with arrows. **F**. Wild *Procephalobus* sp. strain JU2895, with *N. ciargi* infection. A single class of spore size is observed, often clustered inside vesicles as indicated with arrows. Scale bar: 10 μm.

*N*. *major* and *N*. *minor* also displayed two spore size classes. *N. major* formed slightly longer but thinner spores than *N. parisii*. *N. minor* showed however much smaller spores, for each class taken separately (Table 2, 4; Fig 4 C, D). In contrast, *N. homosporus* and *N. ciargi* only have a single class of spore size, with *N. homosporus* spores having an intermediate size (2.00 ± 0.22 μm long, 0.72 ± 0.12 pm wide) and *N. ciargi* spores having a smaller size (1.39 ± 0.20 pm long, 0.59 ± 0.13 pm wide). Spore vesicles were observed more frequently with *N. homosporus* or *N. ciargi* infections than with other *Nematocida* infections (Fig 4 E, F).

**Table 4.** Spore sizes of each nematode-infecting microsporidia species, as determined by Nomarski optics.

| Nematode-infecting<br>Microsporidia | Spore sizes ($\mu\text{m}$) | |
| --- | --- | --- |
|  | Length | Width |
| *Nematocida parisii*<sup>a</sup> | $3.17 \pm 0.22$ | $1.31 \pm 0.15$ |
| | $2.18 \pm 0.15$ | $0.80 \pm 0.08$ |
| *Nematocida displodere*<sup>b</sup> | $2.38 \pm 0.26$ | $1.03 \pm 0.18$ |
| *Nematocida ausubeli* JUm2009 | $2.77 \pm 0.28$ (2.31 ~ 3.26)<sup>c</sup> | $0.94 \pm 0.16$ (0.63 ~ 1.30) |
| | $2.00 \pm 0.36$ (1.36 ~ 2.67) | $0.52 \pm 0.12$ (0.28 ~ 0.80) |
| *Nematocida major* JUm2507 | $3.44 \pm 0.23$ (3.04 ~ 3.79) | $1.20 \pm 0.21$ (0.90 ~ 1.53) |
| | $2.35 \pm 0.23$ (2.02 ~ 2.66) | $0.57 \pm 0.14$ (0.41 ~ 0.77) |
| *Nematocida major* JUm2747 | $3.29 \pm 0.25$ (2.87 ~ 3.69) | $1.20 \pm 0.23$ (0.79 ~ 1.60) |
| | $2.21 \pm 0.26$ (1.85 ~ 2.60) | $0.53 \pm 0.06$ (0.44 ~ 0.63) |
| *Nematocida minor* JUm1510 | $1.93 \pm 0.18$ (1.54 ~ 2.23) | $0.83 \pm 0.13$ (0.51 ~ 1.09) |
| | $1.31 \pm 0.17$ (1.09 ~ 1.73) | $0.55 \pm 0.12$ (0.35 ~ 0.73) |
| *Nematocida homosporus* JUm1504 | $2.07 \pm 0.30$ (1.69 ~ 2.50) | $0.77 \pm 0.12$ (0.56 ~ 0.94) |
| *Nematocida homosporus* NICm516 | $1.95 \pm 0.14$ (1.67 ~ 2.16) | $0.69 \pm 0.11$ (0.56 ~ 0.91) |
| *Nematocida ciargi* JUm2895 | $1.39 \pm 0.20$ (1.54 ~ 2.24) | $0.59 \pm 0.13$ (0.41 ~ 0.84) |
| *Enteropsectra longa* JUm408 | $3.76 \pm 0.38$ (3.15 ~ 4.99) | $0.49 \pm 0.06$ (0.35 ~ 0.68) |
| *Enteropsectra breve* JUm2551 | $1.81 \pm 0.22$ (1.34 ~ 2.14) | $0.66 \pm 0.12$ (0.42 ~ 0.90) |
| *Pancytospora philotis* JUm1505 | $3.46 \pm 0.48$ (2.42 ~ 4.65) | $0.42 \pm 0.06$ (0.25 ~ 0.52) |
| *Pancytospora epiphaga* JUm1396 | $3.71 \pm 0.31$ (2.99 ~ 4.22) | $0.80 \pm 0.09$ (0.56 ~ 0.92) |

<sup>a</sup> Troemel et al. 2008
<sup>b</sup> Luallen et al. 2016
<sup>c</sup> Values of length and width of each microsporidia spore are given with the average ± SD, followed with range of values in the parentheses.

*N. ausubeli* being the most commonly found parasite of *C. elegans* besides *N. parisii,* we further chose to study its lifecycle by electron microscopy. The ultrastructure by electron microscopy and the deduced lifecycle of *N. ausubeli* overall resembled those of *N. parisii,* with possible differences outlined below. High-pressure freezing/freezesubstitution allowed better to visualize lipid membranes compared to room temperature preparation methods. We observed meronts, which are separated from the host cell by a single membrane bilayer, likely pathogen-derived (Fig 5A, B; S1A, B Fig). Their cytoplasm appeared packed with ribosomes. Some meronts displayed an elongated shape and contained several nuclei (Fig 5B, L). The membrane enclosing the meronts appeared to darken progressively and intracellular membrane compartments developed, likely corresponding to the progressive transition to a sporont stage (Fig 5C). We further observed sporogony, whereby individual sporoblasts with a single nucleus are formed, each surrounded by a membrane (Fig 5D, E; S1A, D Fig). We did not observe any nuclear division at this stage (unlike in *Enteropsectra longa,* where they were easily found; see below). We observed progressive stages of sporogenesis, including formation of the anchoring disk, polaroplast membranes, polar tube, posterior vacuole and spore coat (Fig 5E-G; S1C, D, F, G Fig).

**Fig 5:**
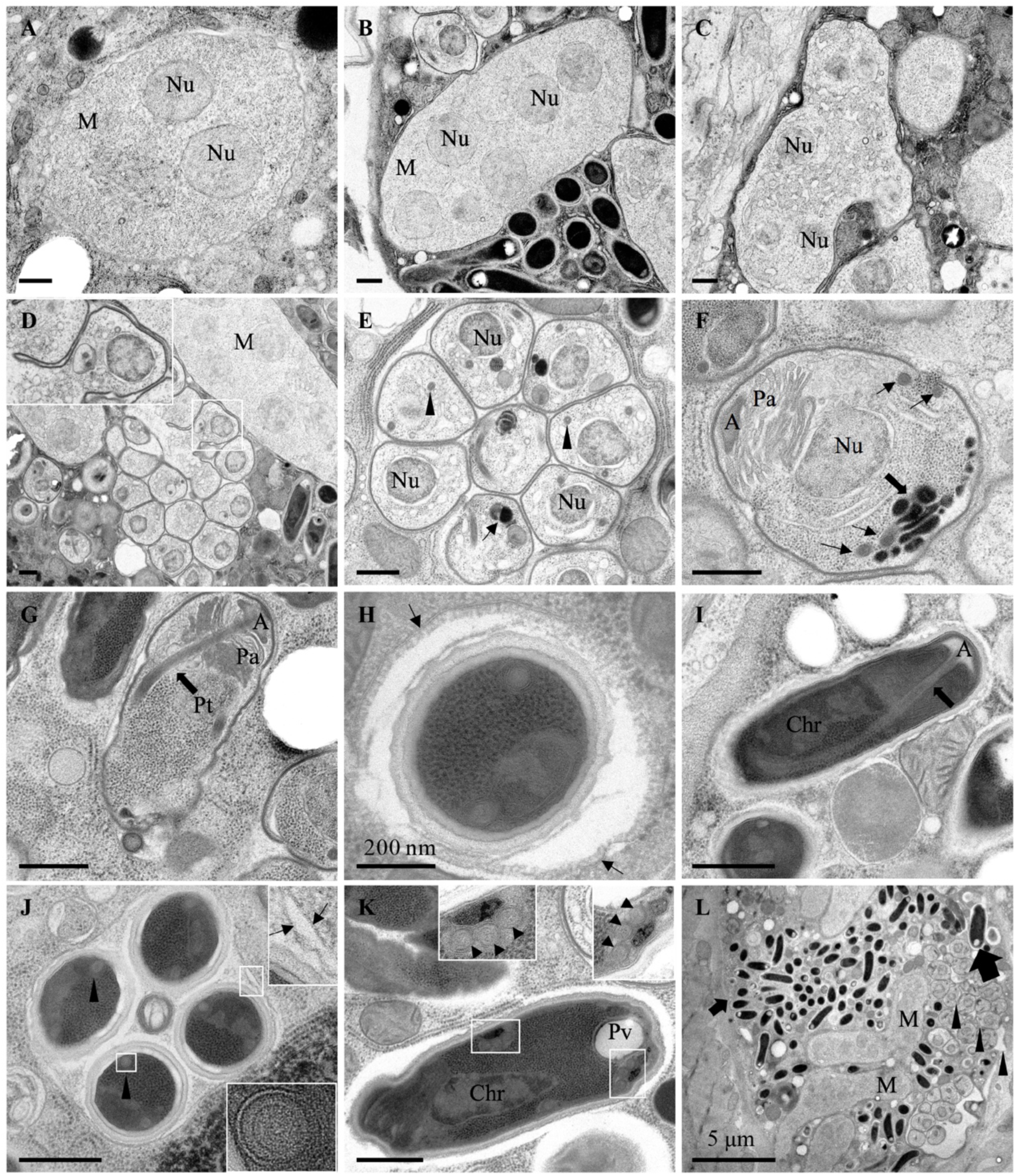
Ultrastructural observations of *Nematocida ausubeli.* Transmission electron micrographs of *N. ausubeli* strain JUm2009 after high-pressure freezing/freeze substitution. **A.** *N. ausubeli* meront with two nuclei. **B.** A multinucleated meront. **C.** Late stage meront. **D.** Formation of sporoblasts by polysporous sporogony. **E.** Cluster of sporonts after sporogony; the arrowheads indicate the nascent polar tube and the arrow indicates the dense membrane structure. **F.** Sporoblast with a maturing anchoring disk and the dense membrane structure on the future posterior side of the spore (large arrow). Four nascent polar tube coil cross¬sections (arrows) are visible, suggesting that this sporoblast may form a spore of large size. **G.** Late stage sporoblast. The arrow indicates the polar tube. **H**. Mature spore with surrounding additional membrane (arrows). The internal side of this membrane is coated. **I.** Mature spore with polar tube indicated by arrow. The anchoring disk and the membranes of the polaroplast are visible on the anterior side, chromatin and ribosomes on the posterior side. **J.** Cross-section of a spore vesicle containing four spores, each showing two polar tube sections (arrowheads). The upper inset shows two membranes around the vesicle (indicated by arrows). The lower inset shows an enlarged multilayered polar tube. **K.** A large size spore, with two insets showing the posterior vacuole and at least three polar tube coils (three cross-sections on either side of the spore, arrowheads). **L.** Lower magnification view of several *N. ausubeli* infection stages in host intestinal cells. Large arrow and small arrow indicate large spore and small spore, respectively. The large spore is that shown in panel K in another plane of section. Arrowheads indicate sporonts. Two multinucleate meronts are indicated. Scale bar is 500 nm, unless indicated otherwise. A, anchoring disk; Chr, chromatin; M, meront; Nu, nucleus; Pa, anterior polaroplast; Pp, posterior polaroplast; Pt, polar tube; Pv, posterior vacuole.

In the final stages of sporogenesis and in mature spores that corresponded to the small size class observed in light microscopy, two polar tube coil cross-sections could usually be observed (Fig 5H, J; S1L Fig). A single large spore could be found, which displayed three polar tube coil sections on either side of the spore (six sections in total; Fig 5K). Thus, the tube coiled several times in large spores, instead of once in the small spores. In *N. parisii,* five polar tube sections were reported on one side of the large spores [22]; it is thus possible that large spores of *N. ausubeli* harbor fewer polar tube coils than those of *N. parisii* (because a single large spore was found in each species, it is however difficult to conclude). The anchoring disk defines the anterior pole of the spore. Below the anchoring disk, the polar tube is lined on either side by polaroplast membranes (visible in Fig 5F, G). A polar tube cross-section with several layers could be seen in Fig 5J and the posterior turn of the polar tube in S1I Fig. The mature spore was seen to contain a posterior vacuole on the side opposite to the anchoring disk (Fig 5K; S1H-J Fig). This vacuole seemed to develop from a dense membrane compartment of the sporoblast (S1C, D Fig). The spores displayed an external coat with several layers (Fig 5H, J, K, 6A; S1I-L Fig).

The spores in the host cytoplasm appeared either isolated, or clustered within a large vesicle. Some isolated spores were surrounded by an additional membrane outside the spore coat and the inner face of this membrane appeared coated (Fig 5H). Unlike in *N. parisii* [27], we could not see the additional membrane around all spores. Fig 6A shows a spore apparently exiting the host cell through exocytosis (although we cannot rule out that such images correspond to endocytotic events). Spores in the lumen were not surrounded by any additional membrane (Fig 5K, L, 6A).

**Fig 6:**
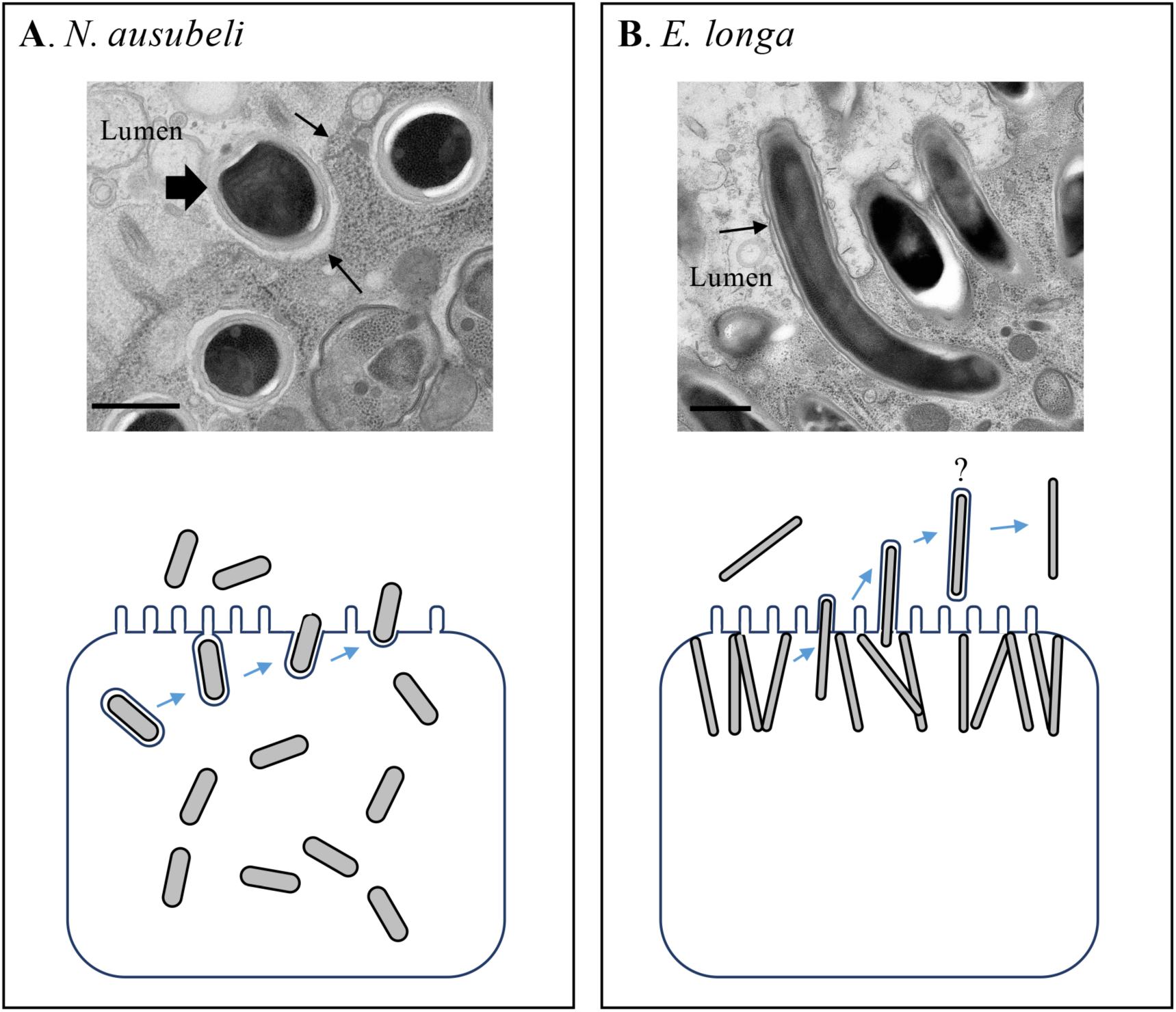
Cell exit modes of *Nematocida ausubeli* and *Enteropsectra longa*. **A.** *Nematocida ausubeli*. The top panel is an electron microscopy image of a *Nematocida ausubeli* spore (large arrow) exiting from the intestinal cell into the lumen. Arrows indicate the apical membrane of the host intestinal cell. The hypothetical diagram below illustrates the exit of *N. ausubeli* spores from intestinal cells by exocytosis. As in **N. parisii** [27,35], spores appear surrounded by a membrane that fuses with the apical membrane of the host intestinal cell, resulting in the release of spores. We also observed apparently mature spores without an additional membrane and do not know whether they will later acquire a membrane or exit in another manner. Earlier stages were omitted here for simplicity. **B.** *Enteropsectra longa*. The top panel is an electron micrograph of *Enteropsectra longa* spores exiting from the intestinal cell into the lumen, with the host intestinal cell membrane folding out around the *E. longa* spores (arrow). The diagram below illustrates the exit of **E. longa** spore from the intestinal cell. The host intestinal cell membrane folds out around the spore until the whole spore exits the cell, after which the host membrane around the spore seems to disappear. Meronts and sporoblasts are not represented in either panel. Scale bars: 500 nm.

When spores were clustered in a vesicle, two membranes could be observed around them (Fig 5J, and other instances).

### Nematode-infecting microsporidia in Clade IV

Whereas the *Nematocida* genus is in Clade II of the microsporidia [22,33], the remaining nine microsporidia strains in our collection were placed in Clade IV, which, unlike Clade II, contains several human-infecting microsporidia (Fig 2). This clade assignment was based on SSU rDNA sequences, which had closest (88–89% identities) to the insect parasite *Orthosomella operophterae* (host: moth *Operophtera brumata)* (Table 2). Only four β-tubulin sequences could be obtained, and these were closest (75% ~ 76% identity) to *Vittaforma corneae,* a human-infecting microsporidia species and a close relative of *Orthosomella operophterae* (whose β-tubulin sequence is not available), consistent with rDNA analysis. We thus isolated nematode-infecting microsporidia that are in a distinct evolutionary branch compared to *Nematocida* and are closer relatives of the human-infecting microsporidia.

Eight out of the nine strains in this group have *Oscheius* species as their nematode host and infect their gut: seven of them from different locations in France were found in *O. tipulae,* while JUm408 was found in *Oscheius* sp. 3 [21] from Iceland. The ninth strain, JUm1396, was isolated from a *C. brenneri* strain and is the only one in this set to infect non-intestinal tissues.

In the phylogenetic analysis, these nine strains separated into two groups, corresponding to the two new genera described below, *Enteropsectra* and *Pancytospora* (see section on Taxonomy) (Fig 2; S4 Fig). The first group included four strains, JUm408, JUm1456, JUm2551 and JUm1483, which were phylogenetically placed as a sister group to *Liebermannia* species (with hosts such as grasshoppers) (Fig 2). In the β-tubulin phylogeny, *Enteropsectra* strains also showed a sister relationship with the group of *V. corneae* and *Enterocytozoon bieneusi*, a human intestinal parasite (S2 Fig). However, with β-tubulin, JUm408 and JUm1483 formed a branch, JUm1456 and JUm2551 another branch, which was different from their SSU rDNA phylogenetic position. Based on molecular sequences, spore morphology and host specificity (below), we describe two species in the *Enteropsectra* genus, *E. longa* (type strain JUm408) and *E. breve* (type strain JUm2551), and do not assign the two other strains to a species. *E. longa* and *E. breve* strains have a small mean SSU genetic distance of 0.005 (Table 3) but differ in spore size and host specificity (see below). While *E. longa* and *E. breve* form a sister group to *Liebermannia* species on the SSU rDNA phylogeny, they have a smaller mean genetic distance to *O. operophterae* (0.08) than to *Liebermannia* (0.11).

The second new clade of nematode-infecting microsporidia includes the five remaining strains and showed strong support as sister lineage to the clade formed by *Enteropsectra* and *Liebermannia* species, with *O. operophterae* as outgroup (Fig 2). Based on molecular sequences, host and tissue specificity, we describe two new species: *Pancytospora philotis* (JUm1505 as type strain, JUm1505, JUm1670, JUm2552), found in the *Oscheius* gut, and *P. epiphaga* (JUm1396) from a *C. brenneri* strain from Colombia caused an epidermis and muscle infection (Fig 7; S5 Fig).

**Fig 7:**
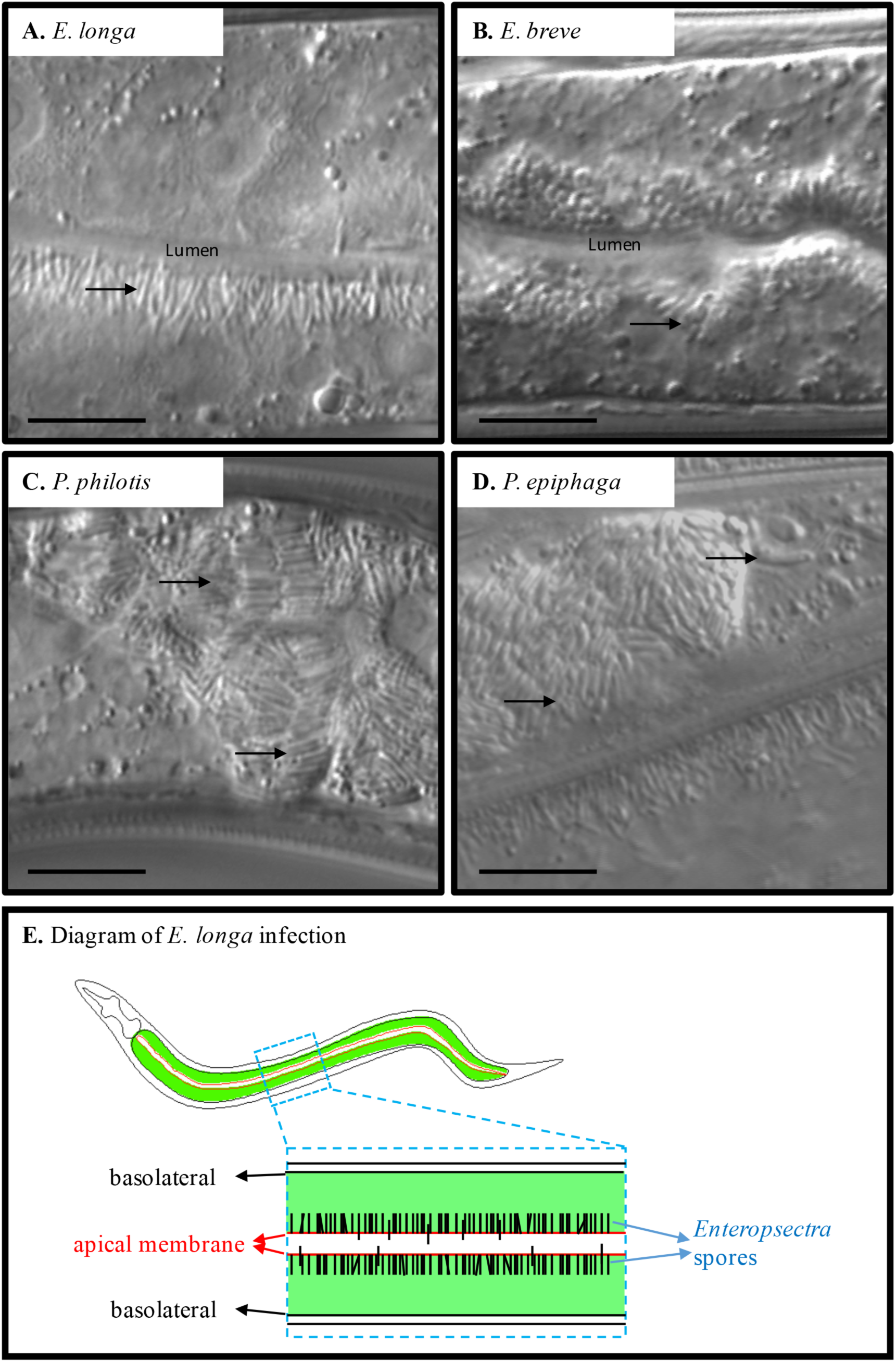
Spore morphology of *Enterospectra* and *Pancytospora* species by Nomarski optics. A. Wild *Oscheius* sp. 3 strain JU408, with *Enteropsectra longa* infection. Gut lumen was indicated. Arrow indicates spores, here long and thin spores that are aligned along the apical side of intestinal cells. **B.** Wild *O***. tipulae** strain JU2551, with *Enteropsectra breve* infection. Arrow indicates spores, here small spores along the apical side of intestinal cells. **C.** Wild *O. tipulae* strain JU1505, with *Pancytospora philotis* infection. Spores are found throughout the intestinal cell. **D**. Wild *Caenorhabditis brenneri* strain JU1396, with *Pancytospora epiphaga* infection. Spores are seen in the epidermal cells in the tail. Scale bar: 10 μm in A-D. **E.** Two-dimensional diagram of *Oscheius* sp. 3 intestine infected with *Enteropsectra longa*. The intestine is formed of polarized epithelial cells. *Enteropsectra longa* starts to form spores along the apical side of the intestinal cells.

### Tissue tropism and lifecycle of nematode-infecting Clade IV microsporidia species

As with *Nematocida,* all of the infections by Clade IV microsporidian strains mentioned above appeared to be transmitted horizontally, as bleaching of the nematode culture eliminated the infection. The *Enteropsectra* strains and *P. philotis* were only observed to infect the intestine of *Oscheius* nematodes. By contrast, *P. epiphaga* (JUm1396) was found to infect epidermis and muscles of *C. brenneri* (Fig 7D; S5D Fig), thus sharing its tissue tropism with *N. displodere,* although on a different evolutionary branch. *P. epiphaga* could also infect *C. elegans* (N2 reference background) (S7F Fig) and *C. briggsae* (AF16).

A striking feature of *Enteropsectra* strains is their cellular localization within the nematode intestinal cells: *Enteropsectra* were all observed to form their spores on the apical side of the epithelial cell at first, while meront stages could be seen in a more basal position (Fig 7A, B, E; 8L). This polarization within the host intestinal cell was not observed in infections of *P. philotis* nor of any *Nematocida* species (Table 2; Fig 4, 7).

The *Enteropsectra* and *Pancytospora* species displayed quite different sizes and shapes of spores from those of *Nematocida* species and we did not see any spore-containing vesicles in these microsporidian infections. They all show a single class of spore size. Though apart in the phylogenetic analysis, *E. longa* (JUm408) and *P. philotis* share similar dimensions of spores, which are particularly long and thin: *E. longa* (JUm408) spores measure 3.76 ± 0.38 μm by 0.49 ± 0.06 μm, while *P. philotis* spores measure 3.46 ± 0.48 pm long by 0.42 ± 0.06 pm. These spores are even longer than the largest spores and thinner than the smallest spores in *Nematocida*. In stark contrast, *E. breve* (JUm2551) form small rod-shaped and crescent-shaped spores (Fig 7B; Table 4).

Because of the striking difference in spore distribution, we further analyzed by electron microscopy the type species of the *Enteropsectra* genus, *Enteropsectra longa* (JUm408) in *Oscheius* sp. 3 JU408. The meront stage appeared overall similar to that of *Nematocida* species: the early stages displayed a cytoplasm packed with ribosomes and very few membranes (Fig 8A); elongated multinucleated meronts could also be observed (Fig 8B). The parasite membrane then progressively darkened, indicating the transition to the sporont stage (Fig 8C-F). Figures of intranuclear mitosis could be seen at this stage, with intranuclear microtubules and spindle plaques at the nuclear membrane (Fig 8E; S6A Fig). Signs of sporogenesis then developed, with a nascent polar tube (Fig 8F-H; S6B Fig). The spore membrane and nascent wall appeared wrinkled (Fig 8G) before becoming smooth in mature spores (Fig 8H-J). The spore wall with its endospore and exospore layers could be clearly observed (Fig 8J). An anchoring disk formed (Fig 8I,K), but the polaroplast membranes were less developed than in *Nematocida* species. In most spores, the polar tube presented a single section (Fig 8G, H, J; S6D Fig). The polar tube however could be seen to turn on the posterior side of the spore (S6C Fig) and occasionally two polar tube sections could be counted in the same spore section (S6E, F Fig). The polar tube thus likely folds back anteriorly on the posterior side of the spore on a short part of its length.

**Fig 8:**
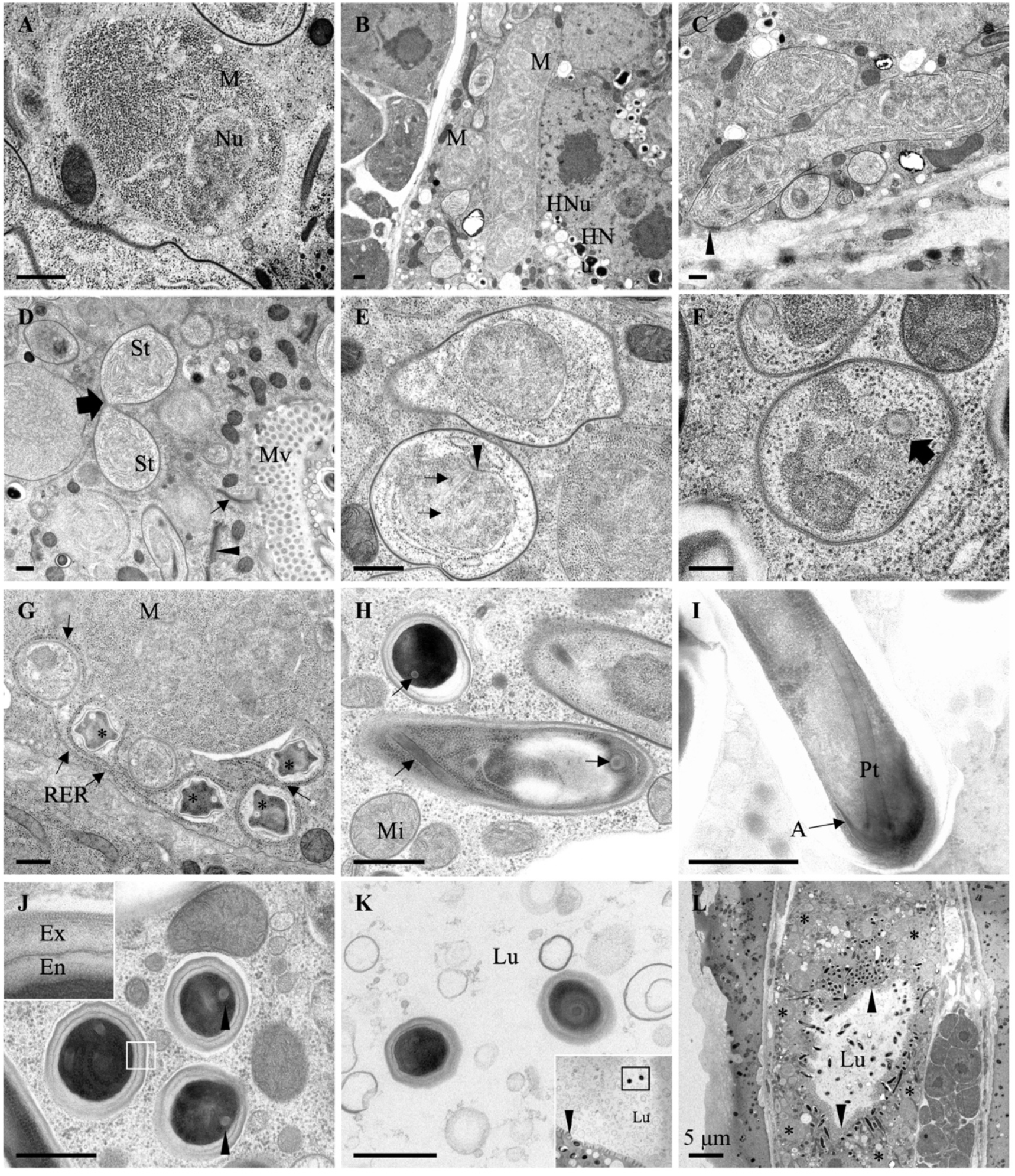
Ultrastructural observations of *Enteropsectra longa.* Transmission electron micrographs of *E. longa* strain JUm408 after high-pressure freezing/freeze substitution. **A.** *E. longa* meront. A nucleus is visible in the cytoplasm full of ribosomes. **B.** Lower magnification with a multinucleated meront. Two meronts are indicated, one with a single nucleus in the plane of section (left) and one with several nuclei (right). Two host nuclei are visible on the right, with a dark nucleolus. Intestinal cells contain two nuclei. **C.** Early sporonts with an electron-dense coat indicated by arrowhead. **D.** Sporont undergoing a cell division (big arrow); small arrow indicates junction of host intestinal cells; the arrowhead indicates a host Golgi apparatus. **E.** Mitotic spindle (arrows designate microtubules) in a sporont; the spindle plaque is indicated by an arrowhead. **F.** Nascent polar tube (arrow) in a sporoblast. **G.** Wrinkled sporoblasts (*). Arrows indicate the host rough endoplastic reticulum folding around the microsporidia. **H.** Late stage sporoblast in the center, mature spore on the top left; arrows indicate polar tubes. **I.** Mature spore with the anterior part of the polar tube, including the anchoring disk. **J.** Cross-section of mature spores. The exospore and endospore layers are shown in the inset. Arrowheads indicate polar tubes. **K.** Two mature spores in the intestinal lumen that do not show an additional membrane around them. Low magnification inset shows the positions of the two spores in the lumen and arrowhead indicates host microvilli. **L.** Low magnification view of cross-section of host, with the intestinal lumen in the center. *E. longa* spores (arrowheads) concentrate around the apical membrane of intestinal cell, while meronts and early sporonts are on the basal side. Scale bar is 500 nm, unless indicated otherwise. A, anchoring disk; Chr: chromatin; Ex, exospore; En, endospore; Lu, lumen; M, meront; Mi: host mitochondrion; Mv, host microvilli; Nu, nucleus; HNu host nucleus; Pt, polar tube; RER, rough endoplastic reticulum; St: sporont.

By electron microscopy, we observed a potential key difference in the exit mode of the spores between *Enteropsectra longa* (JUm408) on one hand, and *N. parisii* and *N. ausubeli* on the other hand. First, the sporoblasts and mature spores of *E. longa* were never seen to be surrounded by an additional membrane outside the spore wall, precluding exocytosis as an exit route. Second, the spores were seen to protrude on the apical side of the host cell, pushing out the host cell membrane like a finger in a glove (Fig 6B; S6G-I Fig). We further focused on spore sections in the intestinal lumen and saw both spores with a surrounding membrane (S6I Fig) and spores without any membrane (Fig 8K).

On the host side, rough endoplasmic reticulum was often seen to wrap around sporoblasts, yet never encircling them fully (Fig 8G). The host cell nuclei presented a characteristic nucleolar structure, which became organized in long tubules (often appearing circular in cross-sections;. S6J, K Fig). On one occasion, microsporidian spores were observed within the host intestinal cell nucleus, whose nucleolus had apparently further degenerated (S6L Fig).

### Host specificity: natural associations and laboratory infection tests

The pattern of natural association revealed an apparent specificity of a given microsporidian species for a nematode genus, mostly *Caenorhabditis* versus *Oscheius* in our collection. Strikingly, *N. parisii, N. ausubeli* and *N. major* infections were found in *Caenorhabditis* species, while *N. minor, N. homosporus* and Clade IV microsporidia species infections were all found in *Oscheius* species and not in *Caenorhabditis* (or, for *N. homosporus,* in *Rhabditella,* a closer relative of *Oscheius* compared to *Caenorhabditis*; Fig 3). The notable exception in Clade IV was the epidermal *P. epiphaga* JUm1396, found in *C. brenneri.* These results suggested a pattern of host-pathogen specificity between nematode and nematode-infecting microsporidia.

We further complemented these natural associations with infections performed in the laboratory. To test for the capacity of a given microsporidia strain to infect a given host, uninfected nematode cultures (cleaned by bleaching) were exposed to microsporidian spores. We used clean spore preparations from seven microsporidian species (see Materials and Methods), namely *N. parisii, N. ausubeli, N. major, N. homosporus, E. longa, E. breve* and *P. philotis.* On the host side, we focused on four nematode species of two genera: *C. elegans*, *C. briggsae*, *O. tipulae* and O. sp. 3, all of which reproduce through self-fertilizing hermaphrodites and facultative males [19,20]. We favored wild strains that had been found naturally infected with microsporidia and were thus not generally resistant to microsporidian infections (Table 5).

**Table 5.** Nematode-infecting microsporidia specificity.

| Species strain | Replicate | *N. parisii*<br>(JUm2816) | *N. ausubeli*<br>(JUm2009) | *N. major*<br>(JUm2747) | *N. homosporus*<br>(NICm516) | *E. longa*<br>(JUm408) | *E. breve*<br>(JUm2551) | *P. philotis*<br>(JUm1505) | *P. epiphaga*<br>(JUm1396) |
| --- | --- | --- | --- | --- | --- | --- | --- | --- | --- |
|  |  | infected/<br>total | infected/<br>total | infected/<br>total | infected/<br>total | infected/<br>total | infected/<br>total | infected/<br>total | infected/<br>total |
| *C. elegans* | 1 | 14/20 | 8/20 | 11/19 | 4/12 | 0/20 | \ | 0/20 | 10/18<sup>a</sup> |
| N2 | 2 | 17/20 | 8/17 | 10/19 | 8/18 | 0/20 | \ | 0/20 | 15/19<sup>b</sup> |
| *C. elegans* | 1 | 10/19 | 10/20 | 14/20 | 2/20 | 0/20 | 0/20 | 0/20 | \ |
| JU2009 | 2 | 11/19 | 5/19 | 5/20 | 5/15 | 0/20 | \ | 0/20 | \ |
| *C. briggsae* | 1 | 10/17 | 6/13 | \ | \ | 0/20 | 0/20 | 0/20 | \ |
| JU2747 | 2 | 14/19 | 7/17 | \ | \ | 0/20 | \ | 0/20 | \ |
| *C. briggsae* | 1 | \ | \ | 13/20 | 4/15 | \ | \ | \ | \ |
| JU2507 | 2 | \ | \ | 12/20 | 9/15 | \ | \ | \ | \ |
| *C. briggsae* | 1 | 11/17 | 11/20 | 12/20 | 9/21 | 0/20 | \ | 0/20 | \ |
| JU2793 | 2 | 12/20 | 13/20 | 14/19 | 10/18 | 0/20 | \ | 0/20 | \ |
| *O. tipulae* | 1 | \ | \ | \ | 13/16 | \ | \ | \ | \ |
| JU1504 | 2 | \ | \ | \ | 13/17 | \ | \ | \ | \ |
| *O. tipulae* | 1 | \ | \ | \ | \ | \ | 6/14 | 11/20 | \ |
| JU1505 | 2 | \ | \ | \ | \ | \ | \ | 7/20 | \ |
| *O. tipulae* | 1 | 0/20 | 0/20 | 0/20 | 0/17 | 0/20 | 4/17 | 3/20 | \ |
| JU1483 | 2 | 0/20 | 0/20 | 0/20 | 1/14 | 0/20 | 5/15 | 8/17 | \ |
| *O. tipulae* | 1 | 0/20 | 0/20 | 0/20 | 1/14 | \ | \ | \ | \ |
| JU170 | 2 | 0/20 | 0/20 | 0/20 | 4/20 | \ | \ | \ | \ |
| *O. tipulae* | 1 | \ | \ | \ | \ | 0/20 | 9/19 | \ | \ |
| JU2551 | 2 | \ | \ | \ | \ | 0/20 | 7/15 | \ | \ |
| *O. sp. 3* | 1 | 0/20 | 8/17 | 0/20 | 9/20 | 11/17 | 3/18 | 0/20 | \ |
| JU408 | 2 | 0/20 | 3/17 | 0/20 | 9/17 | 16/16 | 5/18 | 0/20 | \ |
| *O. sp. 3* | 1 | 0/20 | 8/18 | 0/20 | 11/17 | 12/16 | 3/18 | 0/20 | \ |
| JU75 | 2 | 0/20 | 4/15 | 0/20 | 11/19 | 12/15 | 3/13 | 0/20 | \ |
| *R. typhae* | 1 | \ | \ | \ | 5/12 | \ | \ | \ | \ |
| NIC516 | 2 | \ | \ | \ | 3/13 | \ | \ | \ | \ |

Experiments in the same column are in parallel. The same batch of spores was used on the four nematode species, each with two different strains and two replicates each. At 72 hours post inoculation (hpi) or 120 hpi at 23°C, nematodes found with meronts and/or spores in their intestinal cells were considered infected; otherwise, they were considered uninfected.
<sup>a</sup> Infection were checked at 72 hpi, out of 18 worms, 10 were found infected with meronts, but not spores.
<sup>b</sup> Infection were checked at 96 hpi, out of 19 worms, 15 were found infected with meronts and spores.

*N. parisii* (JUm2816) infected more than 50% of *C. elegans* (N2, JU2009) and *C. briggsae* (JU2747, JU2793) individual animals at 72 hpi. However, no microsporidian infection symptom was observed in *O. tipulae* (JU1483, JU170) nor *O.* sp. 3 (JU408, JU75) at 72 and 120 hpi (Table 5). *O. tipulae* strains JU1504, JU1510 and JU2552 were also exposed to *N. parisii* spores, and none of them became infected either. These infection results indicated that *N. parisii* was unable to infect *O. tipulae* nor *O*. sp. 3 (Table 5).

Specificity of *N. ausubeli* (JUm2009) slightly differed from that of *N. parisii.* By 72 hpi, half of all *Caenorhabditis* animals and about 30% of *O*. sp. 3 showed signs of infection. None of *O. tipulae* worms were infected even at 120 hpi (Table 5). However, when we made a new *N. ausubeli* (JUm2009) spore preparation and used it directly for infection tests, *O. tipulae* strains JU1510 and JU2552 could be infected, but the preparation lost its ability to infect *O. tipulae* over storage at −80°C (see Materials and Methods). We conclude that O. *tipulae* was far less susceptible to infection than *C. elegans, C. briggsae* and *O*. sp. 3 to *N. ausubeli* infection.

The host spectrum of *N. major* (JUm2747) was quite similar to that of *N. parisii:* it infected *C. elegans* (N2, JU2009) and *C. briggsae* (JU2507, JU2793), but not *O. tipulae* (JU1483, JU170) nor *O.* sp. 3 (JU408, JU75) (Table 5). *N. homosporus,* however, could infect both *Caenorhabditis* and both *Oscheius* species and thus appeared as the most generalist (Table 5). Yet *O. tipulae* seemed relatively less sensitive than *C. elegans, C. briggsae* and *O*. sp. 3 to *N. homosporus* infection.

*Enteropsectra* spp. and *Pancytospora philotis* showed different and even opposite specificities compared to the four tested *Nematocida* species. Indeed, none could successfully infect any tested *Caenorhabditis* strains at 120 hpi. Within the two *Oscheius* species, specific interactions were further observed. *Enteropsectra longa* (JUm408) only infected *O*. sp. 3 strains (JU408, JU75), but not *O. tipulae* (JU1483, JU2551), while *E. breve* (JUm2551) infected all four *O. tipulae* and *O*. sp. 3 strains (Table 5). *Pancytospora philotis* (JUm1505) only infected *O. tipulae* (JU1483, JU1505), but not *O*. sp. 3 strains (JU408, JU75). Since *O.* sp. 3 is the closest known species to *Oscheius tipulae, E. longa* and *P. philotis* are examples of narrow specialization in the host-parasite interaction. We also found that *C. elegans* N2 could be infected with *P. epiphaga* (JUm1396), showing epidermal and muscle infection (Table 5; S7F Fig).

The spore morphology of a given microsporidian species was maintained in different nematode species, indicating that host genotype does not affect this pathogen phenotype. For instance, *O. tipulae* (JU1510) infected with *N. ausubeli* (JUm2526) displayed two sizes of spores in its intestinal cells as upon *Caenorhabditis* infection by *N. ausubeli* (S7A Fig). *Oscheius* sp. 3 (JU408) infected with *Enteropsectra breve* (JUm2551) formed small rod-shaped or crescent-shaped spores along the apical side of the worms’ intestinal cells, as upon *O. tipulae* infections (S7D Fig). *C. elegans* N2 infected with *P. epiphaga* (JUm1396) formed long and thin spores in the epidermis and muscles, as upon *C. brenneri* infection (S7F Fig).

### *N. ausubeli* elicits a less robust host transcriptional response than other *Nematocida* species, despite establishing a robust infection

Given the capacity of all *Nematocida* species to infect *C. elegans,* we next sought to compare the *C. elegans* response to infection among our newly isolated microsporidia species. *C. elegans* has been shown to induce a broad transcriptional response after infection by *N. parisii* [36]. Among genes that were highly upregulated at all infection timepoints were *C17H1.6* and *F26F2.1,* two genes of unknown function. Two transgenic *C. elegans* strains, ERT54 and ERT72, were generated as transcriptional reporters for these two genes and have been previously shown to be strongly induced in early *N. parisii* and *N. displodere* infection [36]. We tested these reporter strains with our new microsporidia species by placing them onto plates with a culture of infected worms and microsporidian spores, then monitoring GFP expression at different timepoints in the reporter strains, as well as monitoring microsporidian meront and spore formation. As expected, *N. parisii, N. ausubeli, N. major* and *N. homosporus* could all infect these reporter strains, forming meronts and spores, and induce reporter GFP expression. By contrast, *E. longa* and *Enteropsectra* JUm1483 failed to show evidence of proliferative infection and did not robustly induce reporter expression (Fig 9A-B; S2 Table). Most interestingly, while *N. parisii, N. major* or *N. homosporus* consistently induced the GFP reporters, different strains of *N. ausubeli* (JUm2009, ERTm2, ERTm6; Fig 9; S2 Table) did not, although this species did robustly infect and proliferate within the *C. elegans* intestine.

**Fig 9:**
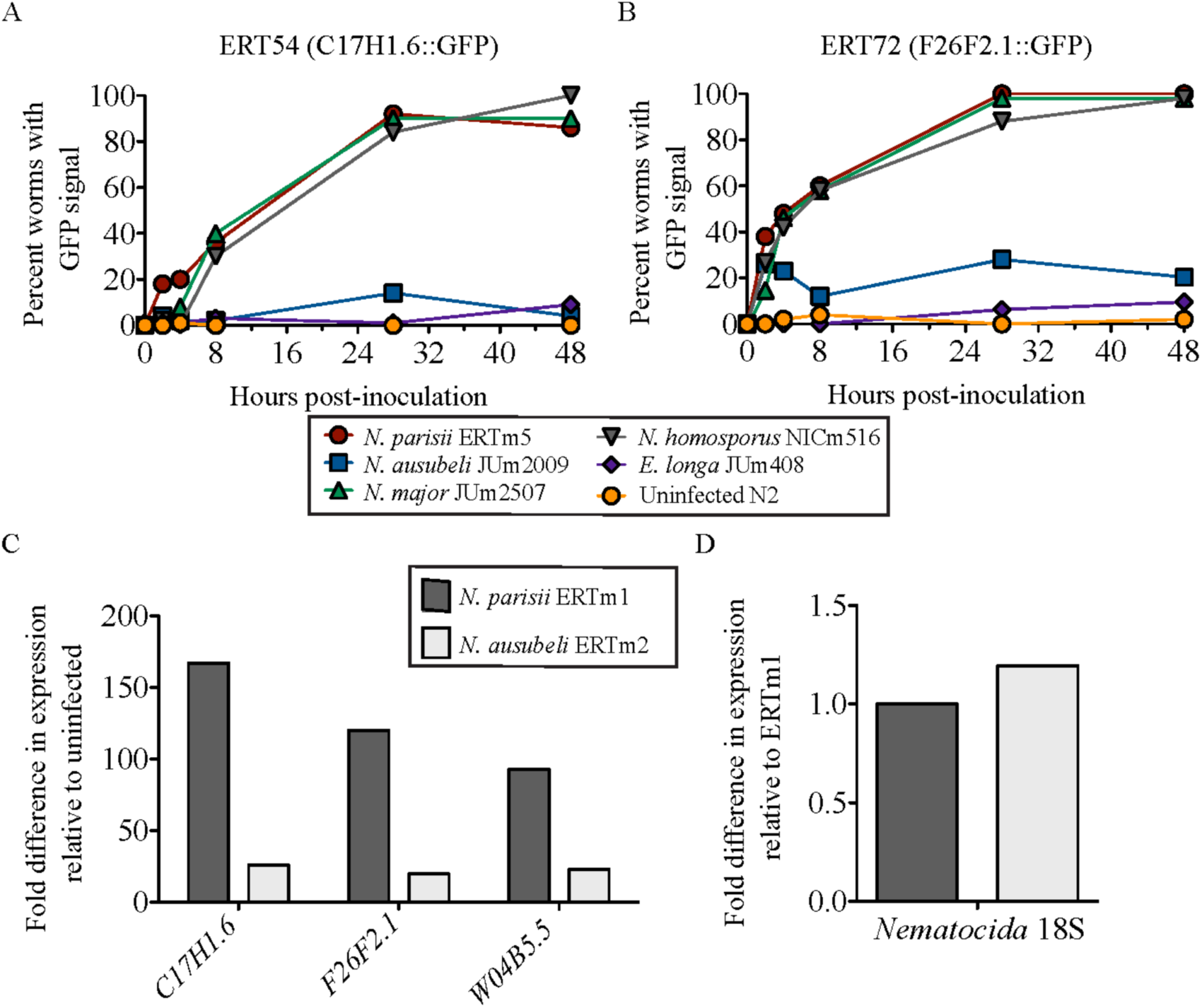
Responses of *C. elegans* strains with transcriptional reporters *C17H1.6p::GFP* and *F26F2.1p::GFP* to exposure by different microsporidia. Strains ERT54 carrying *C17H1.6p::GFP* (**A**) and ERT72 carrying *F26F2.1p::GFP* (**B**) were analyzed for GFP induction at different time points after infection with different microsporidia and the proportion of animals with GFP induction is shown. GFP was reproducibly induced in ERT54 and ERT72 upon infection with *N. parisii*, *N. major* and *N. homosporus,* while GFP signal was rarely observed in ERT54 and ERT72 inoculated with *N. ausubeli* or *E. longa* or the negative control. *N. ausubeli* did infect the *C. elegans* reporter strains, as monitored by DIC as in Table 5. **C**. Transcript levels for three reporter genes were measured after 4 hours of infection of N2 *C. elegans* by *N. parisii* (ERTm1) and *N. ausubeli* (ERTm2). Infection dose was normalized between *Nematocida* by successful invasion events counted as intracellular sporoplasms at 4 hpi. The fold increase in transcript level was measured relative to uninfected N2 levels. **D**. Levels for *Nematocida* SSU rDNA were measured after 4 hours of infection of N2 *C. elegans* by *N. parisii* (ERTm1) and *N. ausubeli* (ERTm2). Infection doses were the same as in panel C and the fold increase in SSU rDNA level was measured relative to N2 infected with ERTm1. The color code indicated on the top applies for panels A and B, the bottom one for panels C and D.

To verify that this differential induction of the GFP reporters matched the transcripts of the endogenous genes, we conducted qRT-PCR after controlled *N. parisii* (ERTm1) and *N. ausubeli* (ERTm2) infections of N2 using purified spore preparations that were normalized for an equivalent level of invasion (see Materials and Methods). Indeed, we saw that both *C17H1.6* and *F26F2.1* transcripts, along with another gene highly induced by *N. parisii* infection *W04B5.5* [37], were induced approximately 6-7-fold lower upon *N. ausubeli* infection compared to *N. parisii* infection (Fig 9C), while the levels of pathogen rRNA (indicative of pathogen load) remained similar (Fig 9D). Thus, N. *ausubeli* infection caused a much reduced host response compared to other *Nematocida* species (Fig 9; S2 Table), despite causing an equivalent, or even more robust infection [38]. Considering the phylogenetic relationships of the *Nematocida* species (Fig 2, 3; S2 Fig), this evolutionary change can be polarized: *N. ausubeli* seems to have lost the capacity to activate these transcriptional reporters as strongly as its closest relatives, or has acquired the capacity to inhibit their activation. Thus, although morphologically quite similar and both are able to infect *C. elegans, N. parisii* and *N. ausubeli* elicit distinct host responses.

## DISCUSSION

### Independent evolutionary branches of nematode parasitism by microsporidia

Microsporidia are ubiquitous obligate intracellular pathogens that have agricultural and medical significance, but have been difficult to study in the laboratory. Our study provides a collection of microsporidia that can infect bacteriovorous nematodes and can easily be studied in the laboratory in their natural hosts and in related species. These rhabditid nematode-infecting microsporidia seem to have more than one origin within the Microsporidia phylum: at least one origin within Clade II and one or two within Clade IV. We thus here enlarge considerably the spectrum of microsporidia that can be cultured in nematodes, including some that are genetically close to human pathogens in Clade IV.

Environmental SSU rDNA microsporidian sequences have been reported from soil, sand and compost samples from North America [39]. (The corresponding species have not been named.) Some of them branch in the SSU phylogeny in the vicinity of the nematode-infecting microsporidia that we isolated (S4 Fig). Specifically, some branch close to *Nematocida homosporus* and some may be outgroups to *Nematocida* or further species of the genus. In Clade IV, one is closely related to the *Pancytospora epiphaga* JUm1396 sequence.

The clades of nematode-infecting microsporidia that we describe have close relatives that infect arthropods, especially insects. This relationship may be due to deep co-evolution (arthropods and nematodes being close relatives on the animal phylogeny), or to the fact that nematodes share their habitats and interact with insects by using them as hosts or carriers [16], which may have facilitated a host shift or a complex lifecycle with several hosts. The microsporidia described here can be cultured continuously in their nematode hosts, but we cannot rule that some of them may use non-nematode hosts as well, including insects. Of note, all of them use a horizontal mode of transmission, despite the fact that many instances of vertical transmission of microsporidia in arthropods, molluscs and fish are known [40,41]. In addition, *Nematocida* species are diploid with evidence of recombination and thus possibly a sexual cycle [29,37], which might occur in another host.

### *N. parisii, N. ausubeli* and *N. major* are relatively common pathogens of *Caenorhabditis* but not of *Oscheius*

Our results suggest that infections by *N. parisii* and *N. ausubeli* are quite common in wild *Caenorhabditis* strains, especially in *C. elegans* and *C. briggsae.* In our collection, *N. parisii, N. ausubeli* and *N. major* infections were found in 30 strains of four *Caenorhabditis* species. Though we have a sampling bias towards France, *N. ausubeli* was found in Asia, Europe and Africa, while *N. parisii* was found mostly in France and once (ERTm5) from Hawaii. *N. major* was only found from three *Caenorhabditis* strains of *C. briggsae* and *C. tropicalis*, all of which were sampled in tropical areas, despite the fact that that we have sampled many hundreds of *C. elegans* isolates and that *N. major* can easily infect *C. elegans* in our specificity infection tests (Table 5). A possibility is that *N. major* may be preferentially distributed in the tropics rather than temperate zones, where *C. elegans* are mostly found (Table 2, Fig 1B) [16].

In addition to *C. elegans* and *C. briggsae* strains, we also have a relatively large collection of microsporidia-infected *Oscheius* strains (10 *O. tipulae* strains and one *O*. sp. 3 strain). However, none of these strains was found with *Nematocida* or *N. major* infections. In line with their natural associations, *N. parisii* and *Nematocida major* were not able to infect any *Oscheius* strains in the laboratory. These specializations may be due to long-term coevolution and adaptation processes [42].

In addition, one new microsporidian species infecting *Caenorhabditis* was found in clade IV, *Pancytospora epiphaga*. As this Clade IV microsporidian species can infect *C. elegans*, it would be interesting to develop its study as a model system for Clade IV species infection.

### Diverse microsporidia infect *Oscheius* species

Microsporidian species that naturally infect *Oscheius* species are diverse (Fig 10, green entries). *N. minor*, found from two *O. tipulae* strains, forms two distinct sizes of spores, similar to *N. parisii, N. ausubeli* and *N. major. N. homosporus* was found from one *O. tipulae* strain and one *R. typhae* and is the only species tested here that is able to infect species of three genera *Caenorhabditis*, *Oscheius* and *Rhabditella*, suggesting that *N. homosporus* may be a relatively less specific pathogen for rhabditid nematodes.

**Figure 10:**
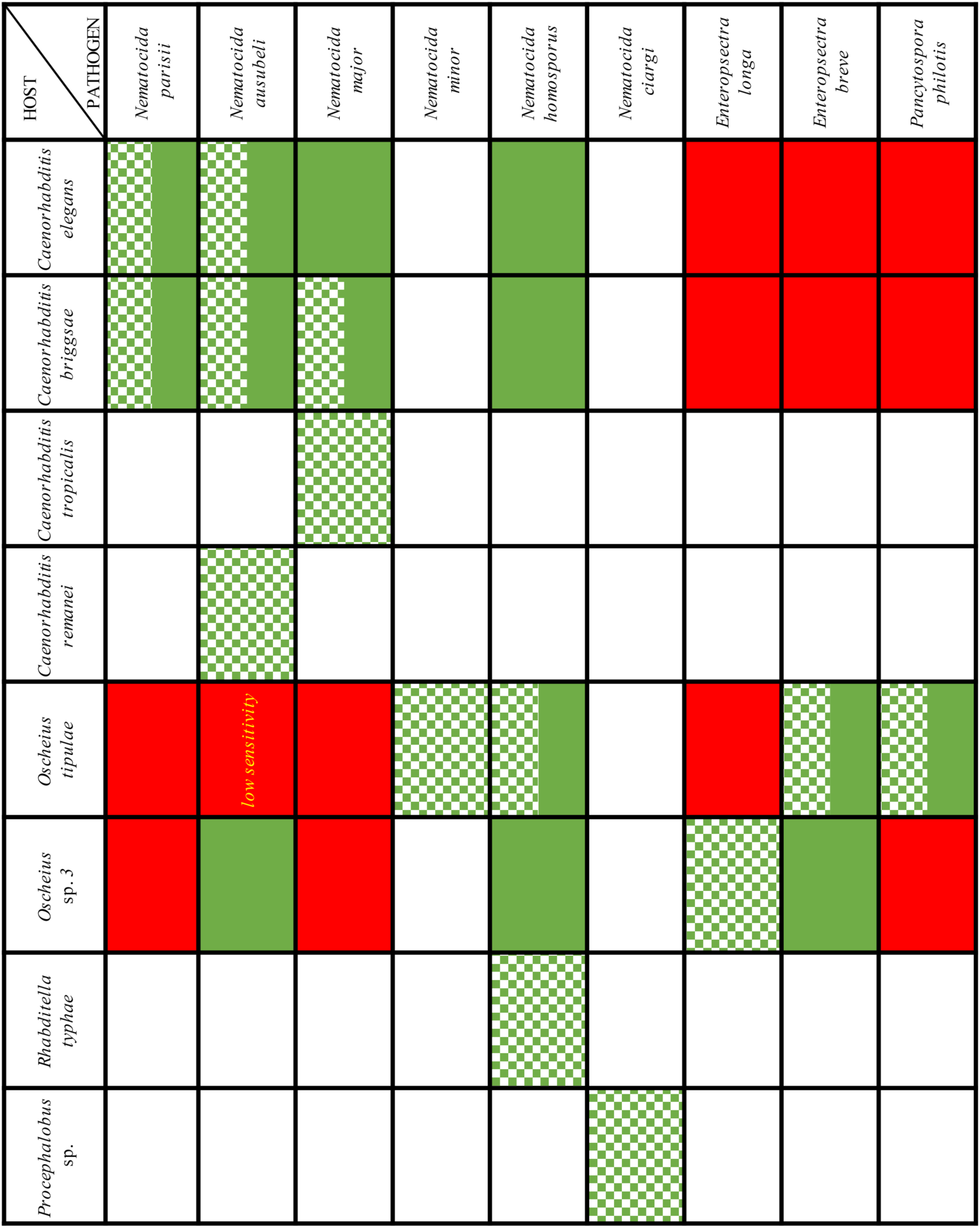
Summary of the interactions between rhabditid nematodes and microsporidia in the wild and in laboratory. A mosaic green color means that the corresponding natural infection was found. Plain green means that the infection worked in the laboratory and red means that the infection did not work in the laboratory. White: not determined.

The Clade IV Oscheius-infecting microsporidia are separated into two groups: *Enteropsectra* species, and *Pancytospora philotis.* None of those could infect *Caenorhabditis* and their host specificity is even narrower, distinguishing between *Oscheius tipulae* and its sister species *Oscheius* sp. 3. The SSU rDNA genetic distances between *E. longa* and *E. breve* are quite small and two other closely related *Enteropsectra* strains are also available (Table 2, 3). Overall, *Enteropsectra* and the *Tipulae* group of *Oscheius* species [19,21] provide an interesting case to study the evolution of a narrow host specificity.

### Evolutionary changes in tissue tropism

Although microsporidia are known to be able to adopt either a horizontal or a vertical transmission [40,43], we here only observed infection in somatic tissues and transmission was horizontal. Most of the infections occurred in host intestinal cells, while two independent instances showed infections elsewhere. As reported previously, *Nematocida displodere* can infect many tissues and cells in *C. elegans*, including the epidermis, muscle, coelomocytes and neurons, although it appears to invade all cells by firing its polar tube from the intestinal lumen [23]. The second independent case is *Pancytospora epiphaga* can be seen in the epidermis, coelomocytes and muscles. Whether it also enters the nematode's cells through the gut remains to be studied.

### Cellular exit strategies

The most striking variation we observed concerns the cellular exit strategies of the spores (Fig 6). *Nematocida parisii* spores acquire an additional membrane around the spore wall and thus exit through a vesicular pathway, using the host exocytosis machinery [27]; in addition, clusters of spores with two additional membranes were observed. If the process is similar in *N. ausubeli* to that in *N. parisii,* the spore clusters may correspond to re-endocytosis of spores from the lumen [35] or perhaps to autophagy of internal spores using the apical plasma membrane. Of note, the host rough endoplasmic reticulum could often be seen to form concentric patterns in the intestinal cell cytoplasm (S1B, E Fig), sometimes wrapping around the sporoblasts (Fig 5E). Whether the reticulum may be a precursor for the additional membranes through an autophagic pathway [44,45], is an alternative possibility.

By contrast, in *Enteropsectra longa,* the sporoblasts and mature spores were never seen surrounded by an additional membrane, which rules out exocytosis as an exit route. Instead, the spores pushed out and deformed the apical plasma membrane of the host intestinal cell (Fig 6; S6 Fig). Whether the final release step was by pinching of the plasma membrane at the base or by rupturing it is unclear, although the former is more probable, given that the intestinal cells were not seen to leak out. We observed spore sections in the lumen, far from any intestinal cells in the corresponding section, with either an additional membrane around them or none. A possible scenario is that the spores are first released with a membrane, and that the membrane then disintegrates (Fig 6B). Yet because we did not follow by serial sectioning the length of the spores, we cannot know for sure that those with a membrane were not still attached to the epithelial cell. We thus cannot rule out an alternative mechanism whereby the spores are released through a hole in the plasma membrane - although given spore size, this latter exit mechanism would likely lead to host cell rupture and death, an event that was never observed.

### A diverse collection of natural host/microsporidia pairs in *C. elegans* and related wild-caught nematodes

Beyond access to a diversity of microsporidia, our collection of host-parasite combinations also provides a resource for defining the genetic basis of host resistance. Most current work on *C. elegans* and *N. parisii* is performed using the *C. elegans* reference strain N2 and the *N. parisii* ERTm1 isolate, yet this strain combination has been shown to lead to a very strong infection where the host does not mount an effective defense response (e.g. in comparison with *C. elegans* CB4856; [28]), thus making it difficult system in which to identify immune defense pathways. The present collection offers many further possibilities of genetic screens using induced mutations or natural genetic variation for resistance pathways.

### Conclusion

Overall, we here considerably enlarged the resources and knowledge on the microsporidia infecting bacteriovorous terrestrial nematodes. These microsporidia are diverse in terms of phylogenetic relationships, spore size and shape, the presence of vesicles containing spores, host specificity pattern, host tissue tropism, host cell intracellular localization and cellular exit route.

## MATERIALS AND METHODS

### Nematode sampling, isolation and microsporidia strains

Hundreds of samples, mostly from rotting fruits, rotting stems and compost, were collected worldwide over several years, and nematodes were isolated as described [11]. The nematode species was identified as described [11,32], using a combination of morphological examination (dissecting microscope and Nomarski optics), molecular identification (18S, 28S or ITS rDNA) and mating tests by crossing with close relatives. Isogenic nematode strains were established by selfing of hermaphrodites or for obligate male-female species from a single mated female. Individuals of strains showing a paler intestinal coloration (Fig 1A) were examined by Nomarski optics. Strains with meronts and spores in the intestinal cells or elsewhere were labeled as suspected to harbor a microsporidian infection. Each nematode strain was then frozen and stored at −80 °C.

For this study, these frozen nematodes were thawed and maintained on nematode growth media (NGM) seeded with *E. coli* OP50 at 23°C. The microsporidian strain was identified after the strain identifier of its host nematode strain (itself identified according to *C. elegans* community rules; http://www.wormbase.org/about/userguide/nomenclature), with an additional "m" between the letters and the numbers for the microsporidia. For instance, a microsporidian strain from the nematode strain JU1762 was named JUm1762. Previously published nematode-infecting microsporidian strains keep their names: ERTm1 (from strain CPA24), ERTm2 (from JU1348), ERTm3 (from JU1247), ERTm4 (from JU1395), ERTm5 (from JU2055) and ERTm6 (from JU1638) [26,28,29,37,46] (Table 1, 2). ERTm4 was previously reported to correspond to *N. parisii* infection [28], but as no sequence data was available in GenBank, we also sequenced the SSU rDNA and β-tubulin genes for this study.

### SSU rDNA and β-tubulin gene sequencing

We sought to amplify by PCR and sequence fragments of two microsporidian genes (SSU rDNA and β-tubulin genes) from all potentially infected rhabditid isolates. Ten infected worms were placed in a PCR tube with 10 μl single worm lysis buffer (1X PCR buffer (DreamTaq Buffer 10X, Theromo Fisher), 1 mM MgCl_2_, 0.45% Tween 20 and 170 ng/μl proteinase K), which was then treated at 60 °C for 60 min, followed by 95 °C for 15 min. This DNA extract was then used as DNA template. To amplify microsporidian SSU rDNA, primers v1f (5'-CACCAGGTTGATTCTGCCTGAC-3') and 1492r (5'-GGTTACCTTGTTACGACTT -3') (Weissli 1994) were used to amplify strains JUm408, JUm1254, JUm1483, JUm1504, JUm2009, JUm2106, JUm2131, JUm2132, JUm2287, JUm2520, JUm2526, JUm2551, JUm2586, JUm2590, JUm2671 and NICm516. We used v1f and 18SR1492 (5'-GGAAACCTTGTTACGACTT-3') to amplify sequences of JUm1456, JUm1460, JUm1505, JUm1510, JUm1670, JUm2552, JUm2793, JUm2796, JUm2799, JUm2816, JUm2825 and JUm2895. We designed a new pair of primers SPF (5'-GATACGAGGAATTGGGGTTTG-3') and SPR (5'-GGGTACTGGAAATTCCGTGTT-3') for JUm2507, JUm2747, JUm2751 and JUm2772. We failed to amplify SSU rDNA for JUm1501 and NICm1041.

To amplify the microsporidian β-tubulin gene, newly designed forward primer βn1F (5'-ACAAACAGGNCARTGYGGNAAYCA-3') and reverse primer βn1R (5'-TGCTTCAGTRAAYTCCATYTCRTCCAT-3') were used. To obtain the β-tubulin gene sequence of JUm2551 and JUm1456, nested PCR was performed using first primers βn1F and βn1R thenβnOF (5'-CCGGACAATATCGTCTTTGG-3') and βnOR (5'-CAGCTCCTGAATGCTTGTTG-3') (S1 Table). PCR products showing a positive signal by gel electrophoresis were sequenced on both strands on ABI 3730XL sequencing machines (MWG). SSU rDNA of five additional *N. parisii* strains (JUm1248, JUm1249, JUm1253, JUm1762, JUm1893) were provided by Aurore Dubuffet and Hinrich Schulenburg. The results were analyzed using Geneious v7.1.7 [47] and compared by BLAST with the NCBI database (http://blast.ncbi.nlm.nih.gov/Blast.cgi). Note that some PCR products could not be amplified (S1 Table). Especially, we failed to amplify the β-tubulin gene in several *Oscheius* infections. Both genes fail to amplify for the putative microsporidian infection of the *O. tipulae* strain JU1501 and this infection could not be characterized.

SSU rDNA and β-tubulin gene sequences have been submitted to GenBank under accession numbers KX352724-KX352733, KX360130-KX360167 and KX378155-KX378171 (S1 Table).

### Phylogenetic analysis

SSU rDNA and β-tubulin gene sequences of microsporidia from this study were analyzed with those of other published microsporidian species and fungi *(Rozella* spp. for SSU rDNA, *Basidiobolus ranarum* and *Conidobolus coronatus* for β-tubulin and concatenated sequences of both genes) as outgroups (Fig 2, 3; S2 Fig) [33]. For phylogenetic analysis of SSU rDNA genes, 28 out of 33 sequences obtained from this study were compared with 11 sequenced *Nematocida* isolates (ERTm1, ERTm2, ERTm3, ERTm5, ERTm6, JUm1248, JUm1249, JUm1253, JUm1762, JUm1893, and JUm2807), JUm1396, 60 other microsporidian species chosen from all five major clades of microsporidia [33] and two *Rozella* species [48]. For analysis of β-tubulin genes, only sequences from the six *Nematocida* species (ERTm1, ERTm2, ERTm3, ERTm5, ERTm6 and JUm2807) and 18 other published microsporidian species were available to be compared with our 32 sequences (S1 Table). To phylogenetically analyze both genes together, we concatenated the two genes of stains ERTm1~6, JUm2807, our 30 strains, 10 other microsporidia species and two outgroups *(B. ranarum* and *C. coronatus).* Sequences were aligned using Geneious v7.1.7 with default parameters and further aligned manually and concatenated if available. The alignments were imported to MEGA 6 [49] to estimate the best DNA evolution models and compute mean genetic distances (1000 bootstrap replicates). Bayesian inference phylogenies were constructed using Mesquite v3.04 [50] and MrBayes v3.2.2 [51], with the same DNA models as above [52] and refined by FigTree v1.4.2 (http://tree.bio.ed.ac.uk/software/figtree/).

### High-pressure freezing and transmission electron microscopy

Worms were frozen in M9 buffer [34] supplemented with 20% BSA (Type V) in the 100 μm cavity of an aluminium planchette, Type A (Wohlwend Engineering, Switzerland) with a HPM 010 (BalTec, now Abra Fluid AG, Switzerland). Freeze substitution was performed according to [53] in anhydrous acetone containing 2% OSO4 + 2% H_2_O in a FS 8500 freeze substitution device (RMC, USA). Afterwards samples were embedded stepwise in Epon. To achieve a good infiltration of spores, the infiltration times in pure resin were prolonged for 48 h compared to the published protocol. After heat polymerization thin sections of a nominal thickness of 70 nm were cut with a UC7 microtome (Leica, Austria). Sections were collected on 100 mesh formvar coated cupper grids and poststained with aqueous 4% uranylacetate and Reynold’s lead citrate. Images were taken with a Tecnai G2 (FEI, The Netherlands) at 120 kV and equipped with a US4000 camera (Gatan, USA).

### Spore size measurements

Spore size was measured as described [22]. Briefly, infected nematodes were photographed by Nomarski optics and spores were measured using the Image J software [54]. 20 spores were measured for each spore type.

### Microsporidia spore preparation

For the microsporidian spore preparation, we first tried the methods previously established for *N. parisii* and *N. ausubeli* [22,46]. Because wild nematodes naturally live in habitats with various microbes [16,17], the microsporidia-infected nematode cultures generally originally contained other microbes, such as bacteria, fungi, or even viruses. In order to obtain a relatively pure microsporidian spore preparation, we treated the nematode cultures repeatedly with antibiotics (100 ug/ml gentamycin, 50 ug/ml Ampicillin, 50 ug/ml Kanamycin, 20 ug/ml Tetracycline, and 50 ug/ml streptomycin), monitoring the presence of *non-E. coli* bacteria and fungi on the plate. Nematode strains do not lose the microsporidian infection after antibiotic treatment. After antibiotic treatment, if the appearance of a plate with infected worms looks like those with bleached worms, we considered the plate to be clean and the infected worms were used to extract clean spores. Even though inconspicuous microbes may still be carried over, as we know so far, none of them could prevent the worms from getting infected with microsporidia nor induce similar symptoms as microsporidia.

Antibiotic-cleaned worms without other detectable microbes were harvested in 2-ml microfuge tube and autoclaved silicon carbide beads (1.0 mm, BioSpec Products, Inc.) were added. The tube was then vortexed for 5 min at 2,500 rpm and the lysate of worms filtered through a 5 μm filter (Millipore) to remove large worm debris. Spore concentration was quantified by staining with chitin-staining dye direct yellow 96 (DY96).

This method worked well on *N. major* and N. *homosporus,* but spores of Clade IV species extracted this way could not infect any worms. To prepare infectious spores of these species, we used instead a plastic pestle to crush worms manually, and stored these spore preparations at 4 °C.

*Nematocida* species spore preparations could generally be stored at −80 °C for later infection tests. However, storage at −80°C could affect the infection efficiency of these spore preparations. Indeed, when we made a fresh *N. ausubeli* (JUm2009) spore preparation and used it directly for infection tests, it could infect *O. tipulae* strains JU1510 and JU2552, with meronts and spores found in their intestinal cells at 120hpi. One month later, we used the same batch that had been stored at −80 °C to infect *C. elegans* (N2), *O. tipulae* (JU1483, JU170, JU1510 and JU2552). At 120 hpi, 100% of N2 adult worms were infected, while none of the *O. tipulae* strains became infected. These results suggested that this spore preparation became less infectious after being frozen and stored at −80 °C for one month, which did not compromise infection in *C. elegans* but did compromise infection of *O. tipulae.* For further specificity tests, spore preparations of *N. major, N. homosporus* and Clade IV species were then used within two hours after extraction, without freezing.

### Infection assays

20 uninfected L4 or young adults (i.e. prior to first egg formation) were transferred to a 6 cm NGM plate seeded with *E. coli* OP50. 5 million microsporidian spores in 100 μl distilled water were placed on the *E. coli* lawn. The cultures were then incubated at 23 °C. The infection symptoms of 20 adults were checked by Nomarski optics at 72 hours after inoculation. If no infection symptoms were found at this timepoint, they were scored a second time at 120 hours post-inoculation.

### Assays with reporter strains

Two transgenic *C. elegans* strains, ERT54 *jyIs8[C17H1.6p::gfp; myo-2p::mCherry]* and ERT72 *jyIs15[F26F2.1p::gfp; myo-2::mCherry]* were used in infection assays to test infection specificity and transcriptional response of *C. elegans* to different microsporidian infections. These two lines express a constitutive fluorescent Cherry marker in the pharyngeal muscles and induce GFP upon infection with *N. parisii* [36]. In the first qualitative assay (23 °C), we focused on the ERT54 strain. First, 10 L4 stage animals from seven naturally infected strains (C. *elegans* JU1762 with *N. parisii* infection, *C. elegans* JU1348 with *N. ausubeli, C. briggsae* JU2507 with *N. major, O. tipulae* JU1504 with *N. homosporus, R. typhae* NIC516 with *N. homosporus, O. tipulae* JU1483 with *Enteropsectra, Oscheius* sp. 3 JU408 with *E. longa)* were transferred to new plates and cultured for two days, in order to release microsporidian spores onto the plates. Then 10 L4 stage worms of the ERT54 strain were added onto these plates and onto a clean plate as control. Two days post-inoculation (dpi), a chunk was transferred to new plate to prevent starvation. One day later (3 days dpi), GFP expression of ERT54 animals (visualized using the Cherry reporter in the pharynx) and infection symptoms were scored. 20 worms showing GFP expression (if any, else the Cherry marker was used) were picked and transferred to a new clean plate. GFP expression was monitored on 8 dpi and 14 dpi. In the second quantitative assay (23 °C), first, 10 L4 stage animals from five naturally infected strains *(C. briggsae* JU2055 with *N. parisii* infection, *C. elegans* JU2009 with *N. ausubeli, C. briggsae* JU2507 with *N. major* infection, *R. typhae* NIC516 with *N. homosporus* infection, *Oscheius* sp. 3 JU408 with *E. longa* infection) and uninfected *C. elegans* reference strain N2 (as negative control) were transferred to new plates and cultured for three days. Then 200 L4 stage worms of ERT54 or ERT72 were added. GFP expression of 50 worms (if possible) of reporter strains was monitored at five different timepoints (2 hours post inoculation (hpi), 4 hpi, 8 hpi, 28 hpi, 48 hpi) and infection symptoms were scored at 48 hpi.

### qRT-PCR of reporter transcripts

For measurements of transcripts levels by quantitative RT-PCR (qRT-PCR) (primers used see S2 Table), 3000 synchronized N2 *C. elegans* L1 larvae were infected for 4 hours at 25°C with 5.0 x 10^5^ ERTm1 *(N. parisii)* spores and 1.5 x 10^6^ ERTm2 *(N. ausubeli)* spores. Prior analysis of serial spore dilutions determined that these ERTm1 and ERTm2 spore doses resulted in an average of 1 sporoplasm per L1 larva at 4 hpi at 25°C as measured by FISH to *Nematocida* rRNA. At 24 hpi, animals were harvested and RNA was isolated by extraction with Tri-Reagent and bromochloropropane (BCP) (Molecular Research Center). cDNA was synthesized from 175 ng of RNA with the RETROscript kit (Ambion) and quantified with iQ SYBR Green Supermix (Bio-Rad) on a CFX Connect Real-time PCR Detection System (Bio-Rad). Transcript levels were first normalized to the *C. elegans snb-1* gene within each condition. Then transcript levels between conditions were normalized to uninfected N2 for *C. elegans* transcripts or normalized to ERTm1 rRNA for *Nematocida* rRNA.

## TAXONOMIC SECTION

### Rationale for the description of new microsporidia taxa

We describe here two new genera and nine new species of microsporidia based on rDNA and β-tubulin sequences and phenotypic analyses.

The rDNA (and β-tubulin, when we could amplify it) sequences could be readily grouped in three distinct clades, one including *Nematocida parisii* and many of our strains in microsporidia clade II, and the two other clades in microsporidia clade IV (Fig 2, 3; S2 Fig).

All described microsporidian species infecting nematodes is reviewed in [26]. Previously described species with associated SSU rDNA sequences are *Nematocida parisii* [22], *Nematocida displodere* [23], *Sporanauta perivermis* [24] and *Nematocenator marisprofundi* [25], the two latter infecting marine nematodes. Compared to the species studied here, *S. perivermis* is found in another group of clade IV, while *N. marisprofundi* appears as a distant outgroup [25] (Fig 2). Our strains are thus all distinct from the two latter species. In addition, two species were reported without any associated molecular sequence [26]. The first one, *Thelohania reniformis,* infected the intestine of a parasitic nematode with a single class of spores of a size exceeding in length and/or width any of those we describe [26]. The second species, of an undefined genus (*"Microsporidium" rhabdophilum*), infected the pharyngeal glands, hypodermis and reproductive system of *Oscheius myriophila* [55], and does not match in tissue tropism and spore morphology any of the present species.

The biological species concept cannot be used in describing these microsporidia as their sexual cycle is unknown and thus we cannot test their crossing ability. Microsporidia species have been classically delimited through their morphology and their association with a host. In recent years, DNA sequences have further helped to assess phylogenetic relationships among microsporidia, and to assign strains to a species when morphology was not sufficient [56]. Among our strains, as a first example, two close groups of strains in the *Nematocida* clade correspond to *N. parisii* and N. sp. 1 in [22], respectively. These groups are consistently distinct from each other by molecular analysis of rDNA and β-tubulin genes (Table 3; S3 Table) but do not appear very different from spore size and general morphology ([22], this work). Their molecular distance (0.017) is consistent with molecular distances between microsporidian species and even greater than other examples of interspecific distance for this gene [57]. Their whole genome sequence [37] further shows that the two species are wide apart, with only 62% amino-acid identity between protein orthologs, while strains of the same species are much closer, such as 0.2% difference at the nucleotide level between *N. parisii* ERTm1 and ERTm3 and 1 SNP every 989 bp for *N.* sp. 1 ERTm2 and ERTm6 [29,37] We therefore formally describe here *N.* sp. 1 as a new species and call it *N. ausubeli* n. sp.

Concerning the other strains in the *Nematocida* clade, given their greater molecular distance to each other, we define four other *Nematocida* species that are also distinct from *N. displodere* [23]. In this case, each of them further shows a distinct spore morphology (Table 4; Fig 4). No other described microsporidian species to our knowledge has a similar sequence nor host distribution. We thus describe them below as four new species of *Nematocida,* namely *N. major* n. sp. (two sizes of spores, each slightly larger than the respective class in *N. parisii* and *N. ausubeli), N. minor* n. sp. (two sizes of spores, each smaller than the respective class in *N. parisii* and *N. ausubeli), N. homosporus* n. sp. (a single class of spores) and *N. ciargi* n. sp. (a single class of spores, particularly small), each with their reference strain. The two latter species were not found in *Caenorhabditis* nematodes but in other bacteriovorous terrestrial nematodes. We could not amplify the SSU gene of *Nematocida* "sp. 7" NICm1041 and therefore refrain from formally describing this putative new species.

The remaining strains of microsporidia in our sampling do not belong to clade II but to clade IV. By blast of the rDNA sequence, they are closest to *Orthosomella operophterae*, an insect pathogen, and by phylogenetic analysis they form two clades. One clade includes JUm408, JUm1456, JUm1483 and JUm2551, and is sister to *Liebermannia* species (also arthropod parasites) - but not particularly close in molecular distance (*Orthosomella* is closer). The other clade includes five strains (JUm1505, JUm1460, JUm1670, JUm2552 and JUm1396) and appears as an outgroup to the four strains + *Liebermannia* spp. Based on the host phylum, the molecular distances and the monophyletic clade groupings, we describe here two new genera named *Enteropsectra* n. gen. for the first group of four strains (type JUm408), and *Pancytospora* n. gen. for the second independent clade (type JUm1505).

In *Enteropsectra* n. gen., we isolated four strains. Based on genetic distance (Table 3; S3 Table), spore morphology (Fig 7, 8; S6 Fig; Table 4), and host specificity (Table 5) of JUm408 and JUm2551, we define two species: *E. longa* JUm408 with large spores (type species of the genus) and *E. breve* JUm2551 with small spores. We do not assign a species name to the two other strains (JUm1456 and JUm1483) as their molecular relationship depends on the gene (SSU rDNA versus β-tubulin). For example, JUm1483 show small spores, was found infecting *Oscheius tipulae* and groups with JUm2551 by SSU rDNA, but its β-tubulin sequence is closer to JUm408. We thus prefer to abstain assigning a species name to this strain.

In *Pancytospora* n. gen., we isolated five strains. Based on genetic distance (Table 3; S3 Table), host specificity and tissue tropism (Fig 7; S5 Fig; Table 5), we define two species: *Pancytospora philotis* n. sp. (type species of the genus) infects *Oscheius tipulae* intestine, while *Pancytospora epiphaga* n. sp. was found to infect *Caenorhabditis brenneri* epidermis and muscles.

### Nomenclatural Acts

The electronic edition of this article conforms to the requirements of the amended International Code of Zoological Nomenclature, and hence the new names contained herein are available under that Code from the electronic edition of this article. This published work and the nomenclatural acts it contains have been registered in ZooBank, the online registration system for the ICZN. The ZooBank LSIDs (Life Science Identifiers) can be resolved and the associated information viewed through any standard web browser by appending the LSID to the prefix “ttp://zoobank.org/” The LSID for this publication is: urn:lsid:zoobank.org:pub:0C31D734-FE13-49F9-8318-ADC6714F316E. The electronic edition of this work was published in a journal with an ISSN.

### Taxonomic descriptions

Phylum Microsporidia Balbiani 1882

### *Nematocida ausubeli* n. sp. Zhang & Félix 2016

LSID urn:lsid:zoobank.org:act:6D7E3D0D-3348-4885-BF64-DF1EE1B7EEBA The type strain is ERTm2. Two strains ERTm2 and ERTm6 have been submitted to the American Type Culture Collection (ATCC, https://www.atcc.org) as PRA-371 and PRA-372, respectively. The type host is *Caenorhabditis briggsae* [58,59], strain JU1348, which was isolated from a mixed sample of decaying vegetal matter (rotting fruits, leaf litter, soil, bark, flowers). The type locality is Periyar Natural Preserve, Kerala, India. The species was also found in *C. briggsae* in Germany and Cape Verde, and *Caenorhabditis elegans* and *Caenorhabditis remanei* in Europe. The ribosomal DNA sequence, deposited to Genbank under Accession JH604648. The genome of the reference strain has been sequenced [37] (accession AERB01000000). The spores are ovoid and measure 2.77 x 0.94 μm (ranges 2.31-3.26 x 0.63-1.30) for the large class and 2.00 x 0.52 μm (ranges 1.36-2.67 x 0.28-0.80) for the small class.. Infection is localized to the host intestinal cells. Transmission is horizontal, via the oral-fecal route. The species is named to honor Dr. Frederick Ausubel and his work on innate immunity of *C. elegans*.

### *Nematocida major* n. sp. Zhang & Félix 2016

LSID urn:lsid:zoobank.org:act:4D7C3F14-187A-4AD1-A62F-DA79BED716E0 The type strain is JUm2507. The type material is deposited as a live frozen culture of the infected host at XXX and in the collection of the corresponding author (MAF; http://www.iustbio.com/worms/index.php). The type host is *Caenorhabditis briggsae* [58,59], strain JU2507, isolated from rotting figs. The type locality is Khao Sok National Park, Thailand. The species was also found in Guadeloupe in *C. briggsae* and *Caenorhabditis tropicalis.* The ribosomal DNA sequence, deposited to Genbank under Accession KX360148. The spores are ovoid and measure 3.4 x 1.2 μm (ranges 2.9-3.8 x 0.8-1.6) for the large class and 2.3 x 0.54 μm (ranges 1.8-2.7 x 0.41-0.77) for the small class. Infection is localized to the host intestinal cells. Transmission is horizontal, presumably via the oral-fecal route. The species is named after the large size of its spores.

### *Nematocida minor* n. sp. Zhang & Félix 2016

LSID urn:lsid:zoobank.org:act:646590BC-E5A8-4FD5-9026-96B276A4D159 The type strain is JUm1510. The type material is deposited as a live frozen culture of the infected host at XXX and in the collection of the corresponding author (MAF; http://www.iustbio.com/worms/index.php). The type host is *Oscheius tipulae* [60], strain JU1510, isolated from compost. The type locality is Hluboka nad Vlatavou near Budweis, Czech Republic. The species was also found in *O. tipulae* in Armenia. The ribosomal DNA sequence, deposited to Genbank under Accession KX360147. The spores are ovoid and measure 1.9 x 0.83 μm (ranges 1.5-2.2 x 0.5-1.1) for the large class and 1.3 x 0.55 μm (ranges 1.1-1.7 x 0.35-0.73) for the small class. Infection is localized to the host intestinal cells. Transmission is horizontal, presumably via the oral-fecal route. The species is named after the small size of its spores.

### *Nematocida homosporus* n. sp. Zhang & Félix 2016

LSID urn:lsid:zoobank.org:act:C959C7AD-DC01-4391-8B18-1B5D02F7349B The type strain is JUm1504. The type material is deposited as a live frozen culture of the infected host at XXX and in the collection of the corresponding author (MAF; http://www.iustbio.com/worms/index.php). The type host is *Oscheius tipulae* [60], strain JU1504, isolated from a rotting *Arum* stem. The type locality is Le Blanc (Indre), France. The species was also found in the nematode *Rhabditella typhae* in Portugal. The ribosomal DNA sequence, deposited to Genbank under Accession KX360153. The spores are ovoid and measure 2.0 x 0.72 μm (ranges 1.7-2.7 x 0.56-0.94). Infection is localized to the host intestinal cells. Transmission is horizontal, presumably via the oral-fecal route. The species is named after the single class of spore size that can be observed in the host.

### *Nematocida ciargi* n. sp. Zhang & Félix 2016

LSID urn:lsid:zoobank.org:act:77EF241F-463D-443A-819D-C32B1BC49332 The type strain is JUm2895. The type material is deposited as a live frozen culture of the infected host at XXX and in the collection of the corresponding author (MAF; http://www.iustbio.com/worms/index.php). The type host is *Procephalobus* sp. strain JU2895 (Cephalobina), isolated from rotting *Trachycarpus* palm fruits. The type locality is Barcelona, Spain. The ribosomal DNA sequence, deposited to Genbank under Accession KX360152. The spores are ovoid and 1.4 x 0.59 μm (ranges 1.5-2.2 x 0.41-0.84). Infection is localized to the host intestinal cells. Transmission is horizontal, presumably via the oral-fecal route. The species is named after its type locality, close to the Centre de Regulació Genòmica in Barcelona, Spain.

### *Enteropsectra* n. gen. Zhang & Félix 2016

LSID urn:lsid:zoobank.org:act:33CE2667-0109-44DA-9878-34FAD4A2F96B This is a novel microsporidian lineage within microsporidian clade II (ref), with *Orthosomella, Liebermannia* as the closest relatives, based on SSU rDNA phylogenetic analyses. The type species is *Enteropsectra longa* n. sp. Zhang & Félix 2016. The genus is named *Enteropsectra* (feminine) after the morphological aspect of the spores at the apical side of the intestinal cells of the nematode host, resembling a bottle brush.

### *Enteropsectra longa* n. sp. Zhang & Félix 2016

LSID urn:lsid:zoobank.org:act:FC304EDC-B2CA-4486-BF8F-095BE0B60E45 The type strain is JUm408. The type material is deposited as a live frozen culture of the infected host at XXX and in the collection of the corresponding author (MAF; http://www.iustbio.com/worms/index.php). The type host is *Oscheius* sp. 3 [21], strain JU408, isolated from a soil sample. The type locality is the Botanical garden of Reykjavik, Iceland. The ribosomal DNA sequence, deposited to Genbank under Accession KX360142. The spores have the shape of a long and thin rod, measuring 3.8 x 0.49 μm (ranges 3.1-5.0 x 0.35-0.68). The polar tube makes one turn at the posterior part of the spore; one or two polar tube sections can be seen in transmission electron microscopy when the spore is cut transversally. Infection is observed in the host epidermis and does not affect the intestinal cells. The spores do not seem to be enclosed as groups of spores in a vesicle. Transmission is horizontal, presumably via the oral-fecal route. The species is named after the long size of its spores.

### *Enteropsectra breve* n. sp. Zhang & Félix 2016

LSID urn:lsid:zoobank.org:act:236607CA-8C44-414D-916C-802C7C67600D The type strain is JUm2551. The type material is deposited as a live frozen culture of the infected host at XXX and in the collection of the corresponding author (MAF; http://www.iustbio.com/worms/index.php). The type host is *Oscheius tipulae* [60], strain JU2551, isolated from a rotting apple. The type locality is an apple orchard in Orsay (Essonne), France. The ribosomal DNA sequence, deposited to Genbank under Accession KX360145. The spores are ovoid and measure 1.8 x 0.66 μm (ranges 1.3-2.1 x 0.42-0.90). Infection is observed in the host intestine and does not affect the intestinal cells. Transmission is horizontal, presumably via the oral-fecal route. The species is named after the short size of the spores.

### *Pancytospora* n. gen. Zhang & Félix 2016

LSID urn:lsid:zoobank.org:act:1ADAC856-ED00-49C9-9906-634BAF38B355 This is a novel microsporidian lineage within microsporidian clade IV, with *Orthosomella, Liebermannia* and *Enteropsectra* n. gen. as the closest relatives, based on SSU rDNA phylogenetic analyses. The type species is *Pancytospora hilotis* n. sp. Zhang & Félix 2016. The genus is named *Pancytospora* (feminine) after the distribution of the spores throughout the cells.

### *Pancytospora philotis* n. sp. Zhang & Félix 2016

LSID urn:lsid:zoobank.org:act:5EA01A65-F5F4-4EE6-A1B6-2726F9CE8579 The type strain is JUm1505. The type material is deposited as a live frozen culture of the infected host at XXX and in the collection of the corresponding author (MAF; http://www.iustbio.com/worms/index.php). The type host is *Oscheius tipulae* [60], strain JU1505, isolated from a rotting peach. The type locality is Le Blanc (Indre), France. The species was also found in *O. tipulae* in other locations in France. The ribosomal DNA sequence is deposited to Genbank under Accession KX360131. The spores have the shape of a long and thin rod, measuring 3.5 x 0.42 μm (ranges 2.4-4.7 x 0.25-0.52). Infection is localized to the host intestinal cells. Transmission is horizontal. The species is named after its specificity to *Oscheius tipulae* (abbreviation *Oti).*

### *Pancytospora epiphaga* n. sp. Zhang & Félix 2016

LSID urn:lsid:zoobank.org:act:41573063-B494-41C3-B0A8-2417DFEB3DCC The type strain is JUm1396. The type material is deposited as a live frozen culture of the infected host at XXX and in the collection of the corresponding author (MAF; http://www.justbio.com/worms/index.php). The type host is *Caenorhabditis brenneri.* The type locality is a private garden in the vicinity of Medellin, Colombia. The ribosomal DNA sequence, deposited to Genbank under Accession KX424959. The spores are ovoid and measure 3.71 x 0.80 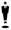 m (ranges 2.99-4.22 x 0.56-0.92). Infection is observed in the host epidermis and does not affect the intestinal cells. Transmission is horizontal. The species is named after the site of infection in the nematode, mostly the epidermis.

## ACKNOWLEDGMENTS

We are very grateful to all sample collectors (M. Asahina, H. Baylis, F. Besnard, A. Zalmanski) and A. Barrière, C. Braendle and L. Frézal for isolating infected strains. We thank M. Botts, K. Balla and M. Bakowski for help with *Nematocida* spore preparations and quantifying reporters and spores; A. Dubuffet and H. Schulenburg for sharing unpublished sequences of *C.* elegans-infecting microsporidia; M. Barkoulas, B. and M. Félix with help with Greek and Latin names, and A. Dubuffet for helpful discussions and comments.

## SUPPORTING INFORMATION CAPTION

**S1-S7 Figure**

**S1 Table. Accession numbers for SSU rDNA and β-tubulin sequences**.

**S2 Table. ERT54 and ERT72 transcriptional reporter induction by various microsporidia**.

**S3 Table. SSU rDNA pairwise distances of all the sequences used for phylogenetic analysis in Fig 2 and genetic distances in Table 3**.

**Supplemental Datafile 1.** Alignment with 116 SSU rDNA sequences used for Fig 2, 3; S6 Fig; Table 3; S3 Table.

**Supplemental Datafile 2.** Alignment with 58 β-tubulin sequences used for Fig 3; S2 Fig. Sequences were aligned using Geneious v7.1.7 with default parameters and further aligned manually and concatenated if available.

